# The paradoxical effects of K^+^ channel gain-of-function are mediated by GABAergic neuron hypoexcitability and hyperconnectivity

**DOI:** 10.1101/2020.03.05.978841

**Authors:** Amy N. Shore, Sophie Colombo, William F. Tobin, Sabrina Petri, Erin R. Cullen, Soledad Dominguez, Christopher D. Bostick, Michael A. Beaumont, Damian Williams, Dion Khodagholy, Mu Yang, Cathleen M. Lutz, Yueqing Peng, Jennifer N. Gelinas, David B. Goldstein, Michael J. Boland, Wayne N. Frankel, Matthew C. Weston

## Abstract

Gain-of-function (GOF) variants in K^+^ channels cause severe childhood epilepsies, but there are no mechanisms to explain how increased K^+^ currents lead to network hyperexcitability. Here, we introduced a human Na^+^-activated K^+^ (K_Na_) channel variant (*KCNT1-Y796H*) into mice and, using a multiplatform approach, found motor cortex hyperexcitability and early-onset seizures, phenotypes strikingly similar to those of human patients. Although the variant increased K_Na_ currents in cortical excitatory and inhibitory neurons, there was a selective increase in the K_Na_ current across subthreshold voltages in inhibitory neurons, particularly in those with non-fast spiking properties, resulting in impaired excitability and AP generation. We further observed evidence of synaptic rewiring associated with hyperexcitable networks, including increases in homotypic synaptic connectivity and the ratio of excitatory-to-inhibitory synaptic input. These findings support inhibitory neuron-specific mechanisms in mediating the epileptogenic effects of K^+^ channel GOF, offering cell-type-specific currents and effects as promising targets for therapeutic intervention.

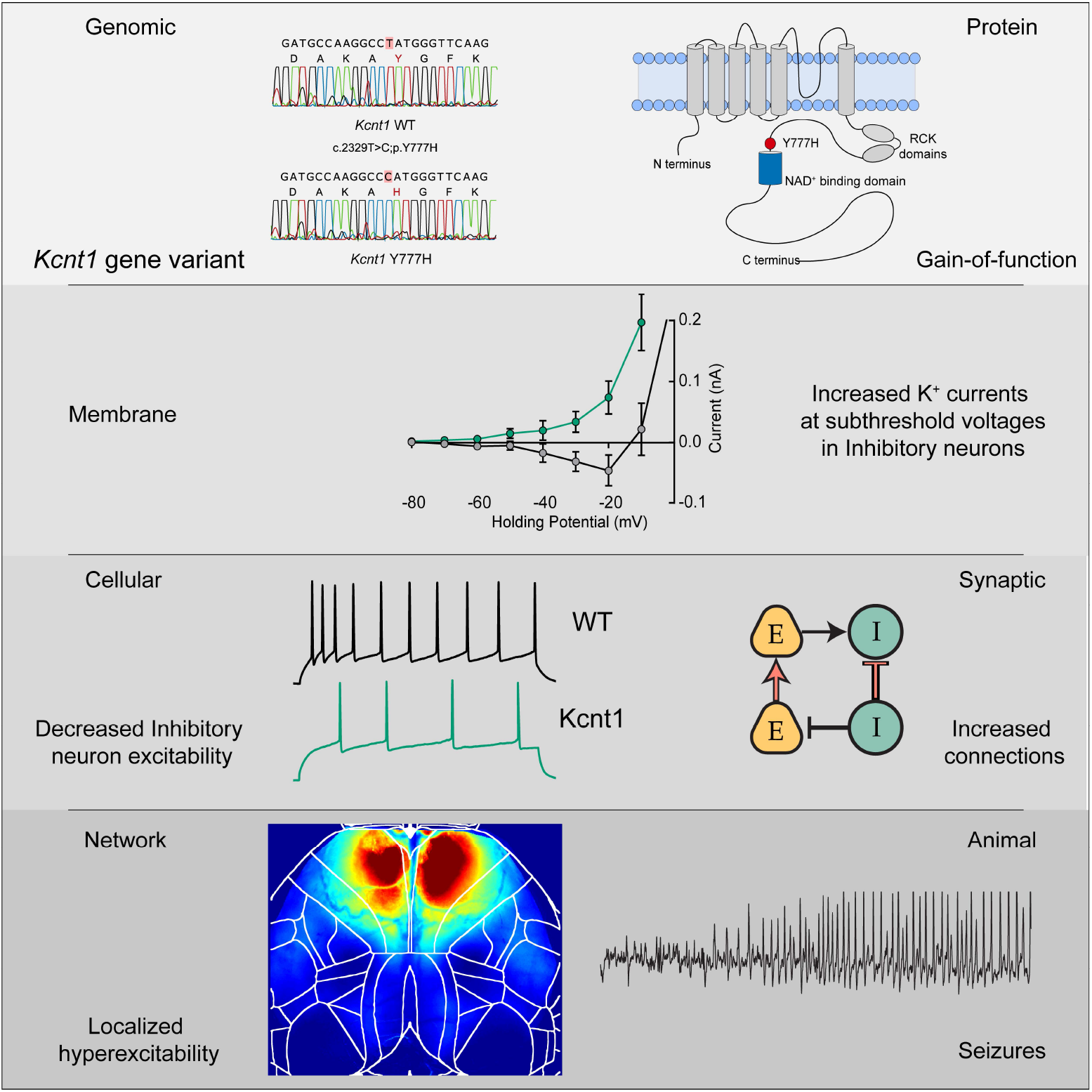
GRAPHICAL ABSTRACT.

## INTRODUCTION

Potassium channels play essential roles in regulating the membrane excitability of neurons, and their dysfunction causes a wide range of neurological diseases, particularly epilepsy (Kohling and Wolfart, 2016, Villa and Combi, 2016, Oyrer et al., 2018). Traditionally, this mechanism has been conceptualized as loss-of-function (LOF) variants decreasing K^+^ currents and leading to neuronal hyperexcitability. Recent human gene discovery efforts, however, have identified novel, neurological disease-associated K^+^ channel variants that increase peak K^+^ current magnitudes when expressed in heterologous cells, suggesting that they are gain-of-function (GOF) variants (Yang et al., 2013, Syrbe et al., 2015, Simons et al., 2015, Millichap et al., 2017, Du et al., 2005, Miceli et al., 2015, Lee et al., 2014). Although these variants have been biophysically characterized, how they function in neurons and networks to cause hyperexcitability is unknown (Niday and Tzingounis, 2018).

Among K^+^ channels, variants in the gene encoding the Na^+^-activated K^+^ channel KCNT1 (Slack, K_Na_ 1.1) cause a range of epilepsies, including early infantile epileptic encephalopathies (EIEE) (Ohba et al., 2015), malignant migrating partial seizures of infancy (MMPSI) (Barcia et al., 2012), and autosomal dominant nocturnal frontal lobe epilepsy (ADNFLE) (Heron et al., 2012, Moller et al., 2015). These variants increase peak K^+^ current magnitude, potentially due to increased channel cooperativity (Kim et al., 2014), enhanced Na^+^ sensitivity, and/or increased channel open probability (Tang et al., 2016, McTague et al., 2018). The broad-spectrum K^+^ channel blocker quinidine can normalize the increased K^+^ currents caused by pathogenic *KCNT1* variants (McTague et al., 2018, Milligan et al., 2014), and has been tried as a precision therapy to treat KCNT1-associated neurodevelopmental disease. Although some early cases were promising (Bearden et al., 2014), further reports show limited therapeutic benefit of quinidine in KCNT1-related epilepsy (Mullen et al., 2018, Chong et al., 2016, Fitzgerald et al., 2019), emphasizing that an understanding of disease mechanisms beyond channel biophysics is necessary to advance clinical treatment.

In normal neuronal physiology, there is evidence that the Na^+^-activated K^+^ (K_Na_) current mediated by KCNT1 can activate in three distinct activity regimes. Either, 1) during bursts of action potentials (APs) to reduce neuronal excitability and control interburst timing (Schwindt et al., 1989, Wallen et al., 2007, Kim and McCormick, 1998, Yang et al., 2007), 2) after a single AP to increase the afterhyperpolarization and enhance bursting (Franceschetti et al., 2003, Liu and Stan Leung, 2004), or 3) at subthreshold voltages to modify intrinsic neuronal excitability (Martinez-Espinosa et al., 2015, Reijntjes et al., 2019). The effect of a KCNT1 GOF variant on neuronal physiology and network activity would therefore depend on which of these regimes were affected. Following this, it has been hypothesized that *KCNT1* GOF variants lead to network hyperexcitability by 1) accelerating AP repolarization selectively in excitatory neurons, thereby limiting sodium channel inactivation and causing a higher AP firing frequency, 2) reducing membrane excitability or inducing spike frequency adaptation selectively in inhibitory neurons, resulting in disinhibition, or 3) causing developmental alterations in synaptic connectivity due to aberrant AP firing (Kim and Kaczmarek, 2014, Niday and Tzingounis, 2018, Milligan et al., 2014). To test these hypotheses, we generated a mouse model with a human disease-causing GOF variant in the *Kcnt1* gene and used a multiplatform approach to identify a time point, brain region, and neuron type that demonstrated strong functional alterations. These results show that the *Kcnt1* variant causes increased currents at subthreshold voltages to modify intrinsic neuronal excitability of non-fast spiking inhibitory neurons and their synaptic connectivity, providing a new paradigm for how GOF variants in K^+^ channels cause neurodevelopmental disease.

## RESULTS

### A mouse model with the *KCNT1* Y796H variant displays epileptic and behavioral phenotypes similar to human patients

The genetic variant Y796H (c.2386T>C; p.[Tyr796His]) in *KCNT1* has been identified as a heterozygous mutation causing both inherited and *de novo* cases of a severe, early-onset form of ADNFLE (Heron et al., 2012, Mikati et al., 2015). To model this *KCNT1*-associated intractable childhood epilepsy in mice, we introduced a missense mutation (c.2329T>C) into the endogenous *Kcnt1* allele on the C57BL/6NJ (B6NJ) strain background (Figure 1A). This mutation encodes a Y777H (p.[Tyr777His]) alteration corresponding to Y796H that lies adjacent to the NAD^+^-binding site (Figure 1B), which is thought to modulate the Na^+^ sensitivity of KCNT1 (Tamsett et al., 2009). We generated cohorts of wildtype, heterozygous, and homozygous littermates (hereafter referred to as WT, *Kcnt1*^m/+^, and *Kcnt1*^m/m^), on the inbred B6NJ background. Genetic transmission of both alleles was normal, and no selective lethality was seen in either mutant genotype at any age. Western blot analysis of membrane and cytosolic protein fractions isolated from the cortices of WT, *Kcnt1*^m/+^, and *Kcnt1*^m/m^ mice showed KCNT1 was only detected in the membrane fraction from WT mice, and importantly, levels and localization of the Y777H variant were unaltered in both *Kcnt1*^m/+^ and *Kcnt1*^m/m^ mice (Figure 1C).

**Figure 1.**
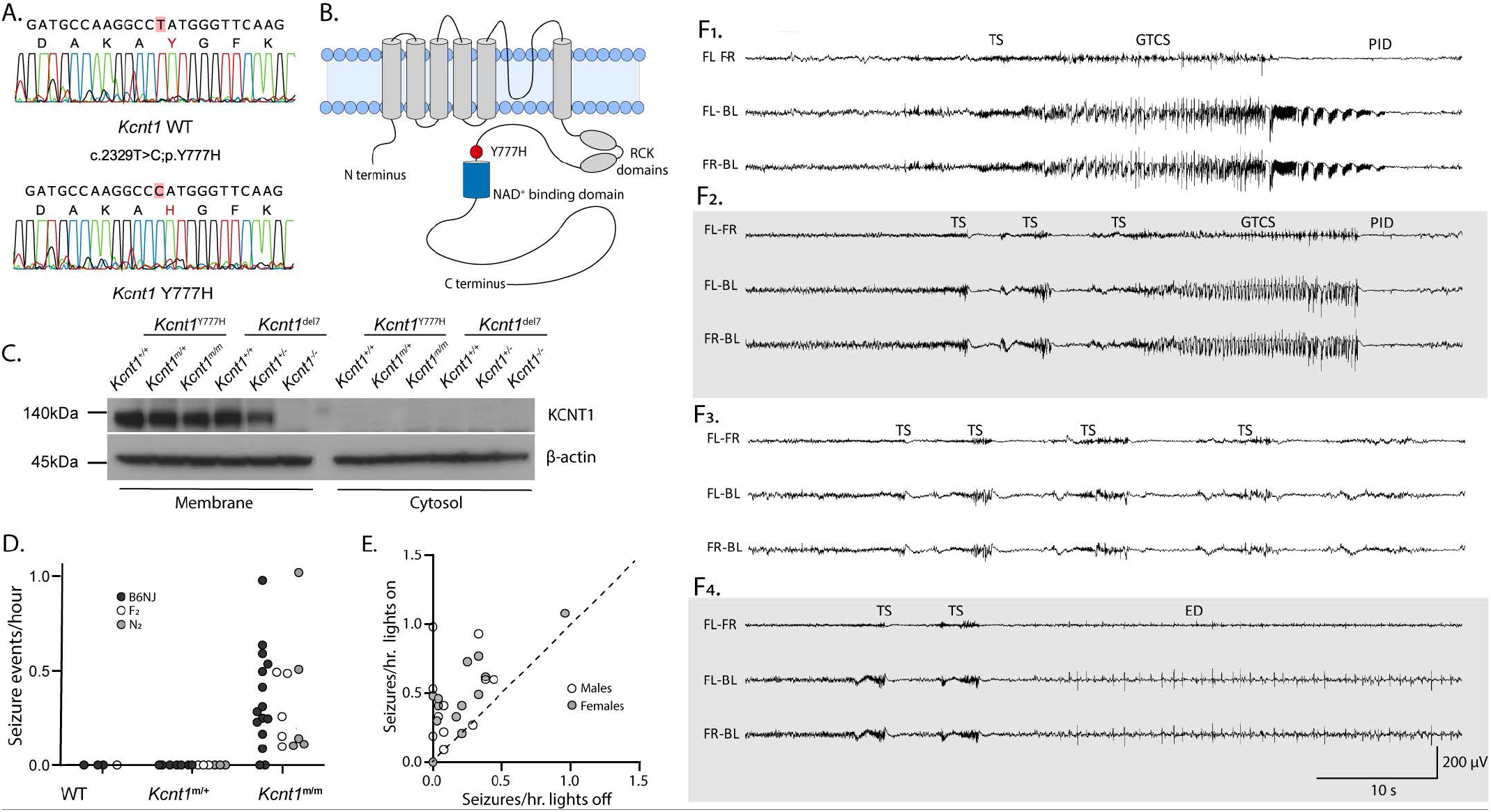
Creation of a *Kcnt1* GOF mouse model in which homozygous mice have two types of spontaneous seizures. **(A)** Sanger sequencing chromatograms show the WT (top) and Y777H (bottom) alleles obtained from the direct sequencing of the PCR products obtained after amplification of the genomic DNA of a WT and a *Kcnt1*^m/m^ mouse. **(B)** Diagram of the KCNT1 protein showing the location of the Y777H residue. **(C)** Western Blots show KCNT1 expression in WT, *Kcnt1*^m/+^, *Kcnt1*^m/m^ and WT, *Kcnt1*^Del7/+^, *Kcnt1*^Del7/Del7^ (a knockout mouse line with a 7bp deletion) P0 cortices following membrane and cytosol fractioning. **(D)** Plot shows the combined spontaneous generalized tonic-clonic seizure (GTCS) and tonic seizure (TS) frequency in WT, *Kcnt1*^m/+^, and *Kcnt1*^m/m^ mice in the three different genetic backgrounds indicated by shaded dots. **(E)** Scatter plot showing that seizures are more likely to occur when lights are on. Each dot represents one *Kcnt1*^m/m^ mouse and is shaded to indicate gender. **(F)** Representative EEG traces showing seizure episodes from B6NJ *Kcnt1*^m/m^ mice, including GTCS, TS, epileptiform discharges (ED), and post-ictal depression (PID) episodes as noted in each panel. The traces shown are differential recordings between front left (FL), front right (FR), and back left (BL) EEG electrodes.

Patients with KCNT1-associated ADNFLE present with frequent, often nocturnal, seizures, mild to moderate intellectual disability, and psychiatric features (Gertler et al., 2018). To test for spontaneous seizures, we monitored adult mice (≥ 7 weeks of age) of each genotype using video-electroencephalography (video-EEG) recordings. Epileptiform activity was not observed in WT (n=3) or *Kcnt1*^m/+^ (n=8) mice. However, 87% (13/15) of *Kcnt1*^m/m^ mice had at least one spontaneous seizure in an average of 38 hours of continuous recording (Figure 1D, black dots). To test the impact of the KCNT1 GOF variant on seizure activity in a broader genetic context, we performed video-EEG on WT, *Kcnt1*^m/+^, and *Kcnt1*^m/m^ mice on a mixed background using an FVB-based partner strain (see Methods). Of the F_2_ and N_2_ hybrid *Kcnt1*^m/m^ mice, 100% (13/13) had at least one spontaneous seizure in an average of 47 hours of continuous recording, whereas seizures were not observed in WT (n=1) or *Kcnt1*^m/+^ (n=6) mice (Figure 1D, gray and white dots). Regardless of mouse background, two distinct seizure types were observed in *Kcnt1*^m/m^ mice: generalized tonic-clonic seizures (GTCS), typically lasting 30-60 seconds, and tonic seizures (TS), typically lasting 1-2 seconds. TS often preceded GTCS (Figures 1F_1_ and 1F_2_), but TS also occurred without further progression, sometimes as single events, but often in clusters of two or more episodes within a few seconds of each other (Figures 1F_3_ and 1F_4_). Because epileptiform activity was not observed in *Kcnt1*-Y777H heterozygous mice, we focused our study on *Kcnt1*-Y777H homozygous mice to understand the contribution of K^+^ channel GOF to seizure disorders.

**Figure 2.**
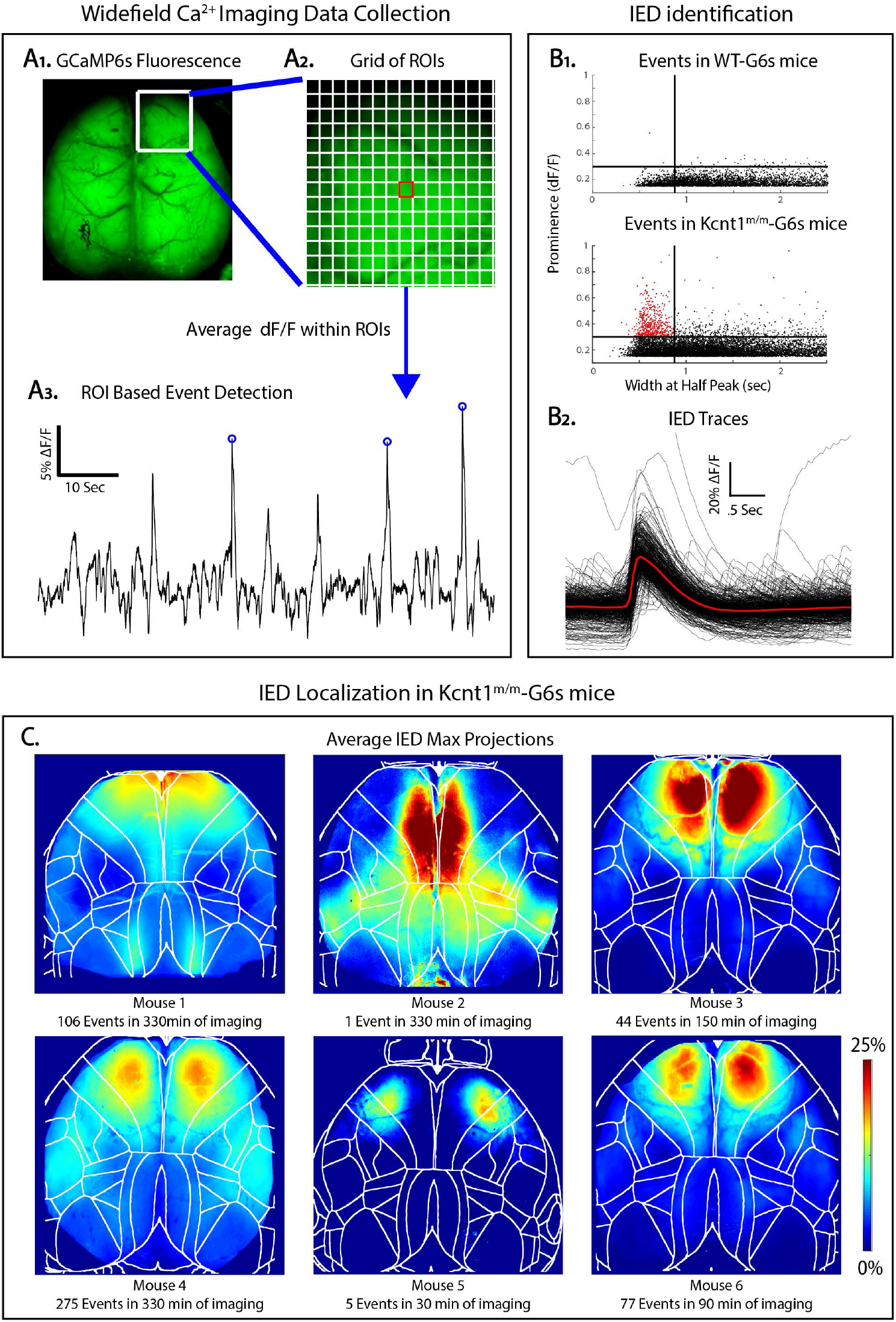
Interictal epileptiform discharges in *Kcnt1*^m/m^ mice localize to the secondary motor cortex. **(A_1_)** GCaMP6s fluorescence in the brain of a *Kcnt1*^m/m^-G6s mouse. **(A_2_)** Illustration of the grid of ROIs used in event detection superimposed on a portion of the GCaMP6s fluorescence image. **(A_3_)** ΔF/F peaks with a prominence greater than 15% were identified in each ROI. **(B_1_)** Maximum event prominence plotted as a function of width at half-max for WT-G6s and *Kcnt1*^m/m^-G6s mice. *Kcnt1*^m/m^-G6s mice generate events (narrow, high prominence) not seen in WT-G6s. Lines at 875 ms and 30% ΔF/F indicate our criteria for classifying events as IEDs. **(B_2_)** Overlayed ΔF/F traces of all IEDs in all *Kcnt1*^m/m^-G6s mice with the mean waveform in red. **(C)** To investigate the cortical localization of spike-like events, we generated a maximum projection across all event frames for each IED. The average of these is shown for each *Kcnt1*^m/m^-G6s mouse, showing consistent peak localized to the anterior medial cortex, mostly within primary and secondary motor cortices. Boundaries indicate cortical regions demarcated by aligning brains to the Allen CCF (See Methods).

**Figure 3.**
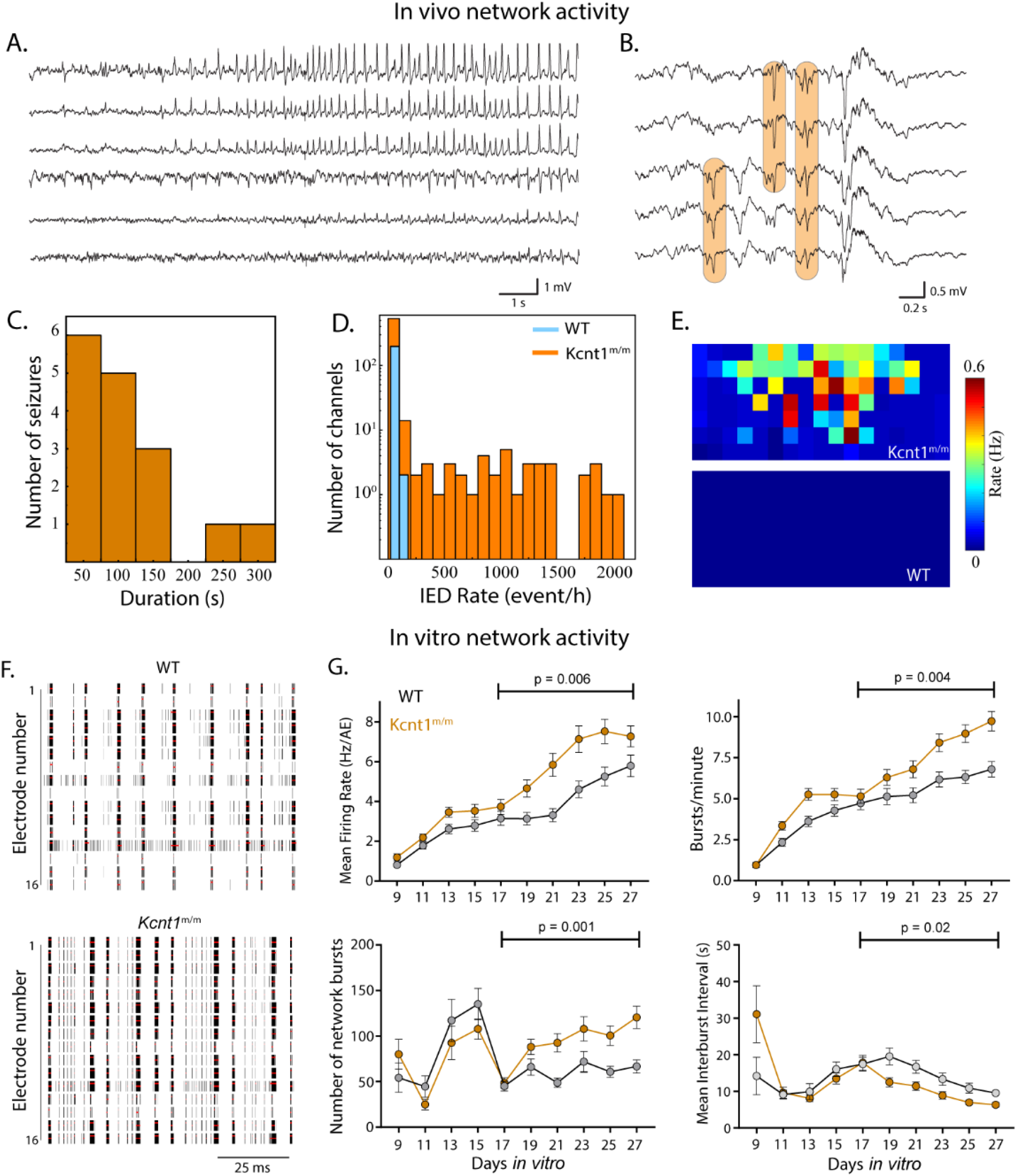
*Kcnt1*^m/m^ mice have epileptic activity *in vivo* at the age of two weeks. **(A)** Sample raw traces from electrocorticography array demonstrating focal, evolving ictal pattern in a head-fixed pup. **(B)** Sample raw traces from surface electrocorticography array demonstrating populations of interictal epileptiform discharges (IEDs; orange boxes). **(C)** Histogram of seizure duration across *Kcnt1*^m/m^ pups. 3/8 *Kcnt1*^m/m^ pups had definite seizures; 0/5 WT pups had seizures. **(D)** Histogram of IED occurrence across all functioning electrocorticography channels in *Kcnt1*^m/m^ (orange) and WT (blue) pups. **(E)** IED occurrence rate localized anatomically across the electrocorticography array and pooled across pups reveals focal distribution in mutants. Each square represents one electrocorticography electrode, and the array is oriented such that anterior brain regions are at the top and medial brain regions are to the right. Warmer colors are associated with higher IED occurrence rate; color map is normalized across *Kcnt1*^m/m^ and WT pups. **(F)** Raster plots showing WT and *Kcnt1*^m/m^ cortical network firing across the 16 electrodes of a well of a representative plate at DIV 23. The black bars indicate spikes, and the red bars indicate bursts. **(G)** Graphs plotting measures of spontaneous activity as a function of days *in vitro* on MEAs of WT (gray, n=65 wells, 5 mice) and *Kcnt1*^m/m^ (orange, n=66 wells, 5 mice) cortical neurons. Mean firing rate (MFR) per active electrodes, number of bursts per minute, number of network bursts, and interburst interval are shown. Permutated p-values for mature DIV 17 to 27 calculated with a Mann-Whitney U test followed by 1,000 permutations are indicated on the graphs.

**Figure 4.**
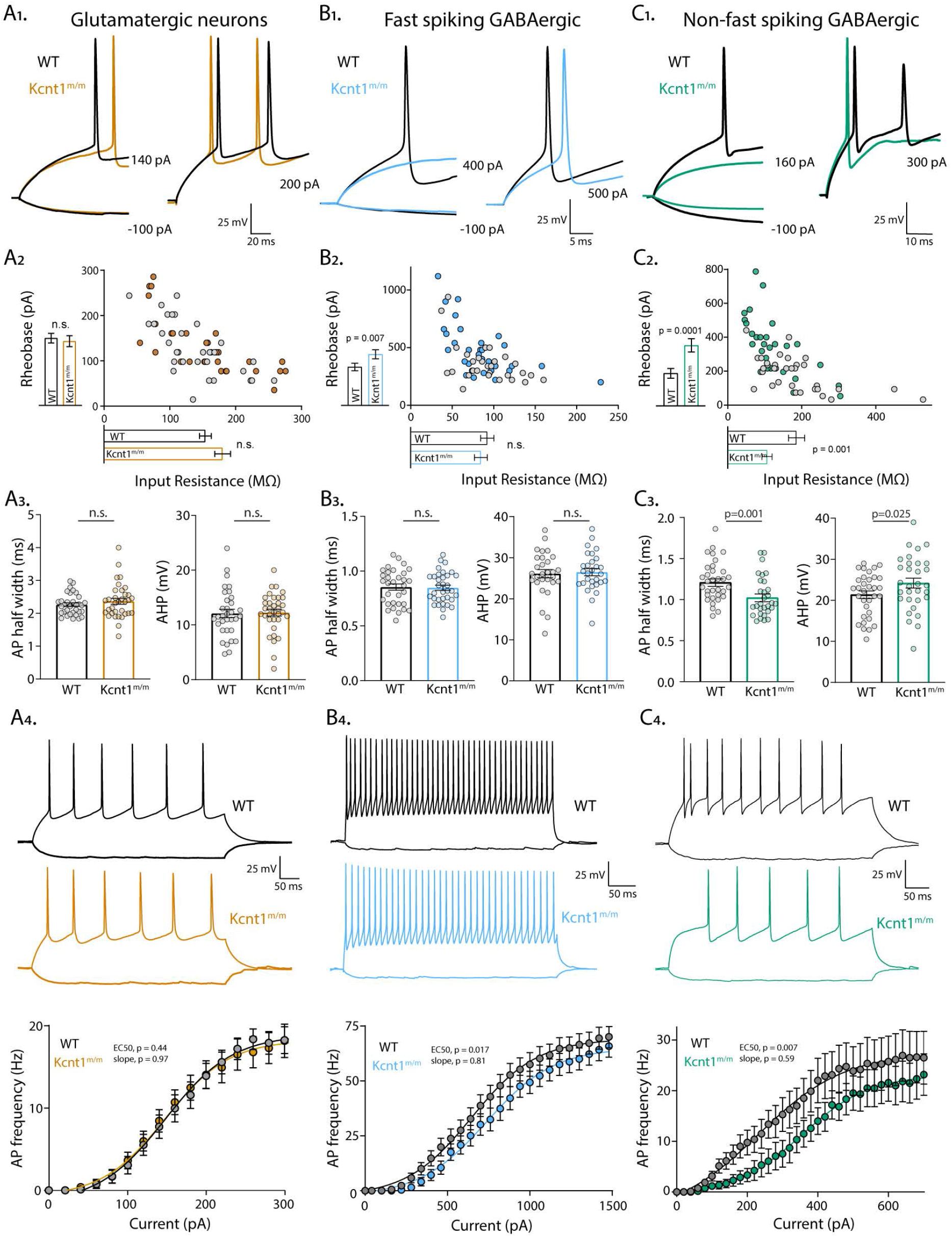
The *Kcnt1* Y777H variant causes a reduction in membrane excitability and AP generation in GABAergic, but not glutamatergic, cortical neurons. **(A_1_)** Representative membrane voltage traces of WT (black) and *Kcnt1*^m/m^ (orange) glutamatergic neurons in response to current injections. **(A_2_)** The individual values of input resistance (x-axis) and rheobase (y-axis) for WT and *Kcnt1*^m/m^ glutamatergic neurons. The mean and s.e.m. for each group are indicated by the bars next to the corresponding axis. **(A_3_)** Individual values and mean ± s.e.m. for AP half-widths and AHP in glutamatergic neurons. **(A_4_)** Example traces and summary data showing the number of APs (mean and s.e.m.) per current injection step in WT and *Kcnt1*^m/m^ glutamatergic neurons. The EC_50_ (WT: 133 ± 9 pA, *Kcnt1*^m/m^: 121 ± 11 pA) and slope (WT: 41 ± 9 pA, *Kcnt1*^m/m^: 42 ± 11 pA) were not different between genotypes. (B_1_) Representative membrane voltage traces of fast spiking GABAergic WT (black) and *Kcnt1*^m/m^ (blue) neurons in response to current injections. (B_2_) Input resistance and rheobase for WT and *Kcnt1*^m/m^ FS GABAergic neurons with mean and s.e.m. indicated by the bars next to the corresponding axis. (B_3_) Individual values and mean ± s.e.m. for AP half-widths and AHP in FS neurons. (B_4_) Traces and summary data showing the number of APs (mean and s.e.m.) per current injection step in WT and *Kcnt1*^m/m^ FS GABAergic neurons. The EC_50_ (WT: 646 ± 24 pA, *Kcnt1*^m/m^: 737 ± 28 pA) was reduced in *Kcnt1* ^m/m^ FS neurons, but not the slope (WT: 198 ± 19 pA, *Kcnt1*^m/m^: 205 ± 21 pA). (C_1_) Representative membrane voltage traces of NFS GABAergic WT (black) and *Kcnt1*^m/m^ (green) neurons in response to current injections. (C_2_) Input resistance and rheobase for WT and *Kcnt1*^m/m^ NFS GABAergic neurons with mean and s.e.m. for indicated by the bars next to the corresponding axis. (C_3_) Individual values and mean ± s.e.m. for AP half-widths and AHP in NFS neurons. (C_4_) Traces and summary data showing the number of APs (mean and s.e.m.) per current injection step in WT and *Kcnt1*^m/m^ NFS GABAergic neurons. The EC_50_ (WT: 231 ± 45 pA, *Kcnt1*^m/m^: 354 ± 23 pA) was reduced in *Kcnt1*^m/m^ NFS neurons, but not the slope (WT: 122 ± 47 pA, *Kcnt1*^m/m^: 96 ± 25 pA). The lines in the current/AP plots are fits to a Boltzman Sigmoidal curve which were used to determine the EC_50_ and slope of the relationship. Statistical significance tested using Generalized Linear Mixed Models.

Across all groups, the incidence of seizure episodes was approximately two-fold higher when the vivarium lights were on, corresponding to the sleep period of mice (Figure 1E). To examine the relationship between seizures and sleep directly, we compared the incidence of seizures in wake, NREM and REM sleep in seven B6NJ *Kcnt1*^m/m^ mice. 90% of seizure episodes (131/145 total) occurred during NREM sleep (Table S1). Thus, independent of mouse strain influence, homozygous expression of the Y777H variant results in frequent, NREM sleep-associated seizures, highlighting meaningful parallels between the mouse model and the human disease caused by the KCNT variant.

Because *KCNT1* GOF variants are associated with intellectual disability and psychiatric conditions, including anxiety and attention deficit hyperactive disorder, we next performed a series of tests to assess the effects of the Y777H variant on mouse behavior. As a global indicator of mouse well-being, we first examined nest building, which is often reduced in neurodegenerative and psychiatric murine disease models (Jirkof, 2014). Both *Kcnt1*^m**/** m^ females and males were poor nest builders compared with WT, as assessed using two different measures - a score of nest quality, and by the weight of unshredded nesting material (Figure S1). In an open field test, *Kcnt1*^m/m^ mice exhibited a robust hyperactivity phenotype, traveling a further distance compared to WT littermates; however, they were not different from WT on the elevated plus maze (Figure S1), indicating hyperactivity without changes in anxiety-like behaviors. *Kcnt1*^m/m^ mice also exhibited decreased freezing behaviors in the contextual and cued fear conditioning tests, without changes in post-shock freezing behavior or acoustic startle responses, suggesting learning deficits un-confounded by sensorimotor deficits (Figure S1). These data suggest that, in addition to recapitulating the seizure phenotypes, *Kcnt1*^m/m^ mice also exhibit several behavioral phenotypes similar to those found in ADNFLE patients.

### Widefield Ca^2+^ imaging reveals network hyperactivity in the secondary motor cortex of *Kcnt1^m/m^* mice

Next, we wanted to know which cortical regions were most strongly impacted by the *Kcnt1* variant. To do this, we crossed *Kcnt1* Y777H mice to the *Snap25*-GCaMP6s line to generate *Kcnt1*^m/m^; *Snap25*^G6s/+^ and *Kcnt1*^+/+^; *Snap25*^G6s/+^ (hereafter referred to as *Kcnt1*^m/m^-G6s and WT-G6s) littermates. We then imaged spontaneous neural activity across the nearly the entire dorsal cortices of awake, head-fixed, behaving *Kcnt1*^m/m^-G6s and WT-G6s adult mice using a custom tandem-lens epifluorescent macroscope (Figure 2A). Both *Kcnt1*^m/m^-G6s and WT-G6s mice exhibited dynamic, spatially complex patterns of spontaneous cortical activity. To quantify this activity, we performed event detection and plotted the width versus prominence profile of all detected events (Figure 2B). In all *Kcnt1*^m**/** m^-G6s mice (n=6), there were narrow, large amplitude calcium events (Figure 2B), similar in appearance to previous reports of interictal epileptiform discharges (IEDs) captured using the same technique (Steinmetz et al., 2017, Rossi et al., 2017). Importantly, these events were never observed in WT-G6s littermates (n=5, Figure 2B_1_). Across *Kcnt1*^m**/** m^-G6s mice, these events were consistently localized to the anterior medial cortex (Figure 2C) consistent with previous reports of IEDs localized to the anterior medial cortex in human patients carrying the Y796H variant (Mikati et al., 2015). Alignment of the activity maps from *Kcnt1*^m**/** m^-G6s mice to the Allen Mouse Common Coordinate Framework (Figures 2C and S2, Allen Mouse Brain Connectivity Atlas, 2017) showed that these events localized primarily to the secondary motor cortex region (M2), suggesting a regional specificity to the interictal hyperexcitability in *Kcnt1*^m**/** m^ mice.

### Cortical network hyperactivity is present at the end of the second postnatal week in *Kcnt1*^m/m^ mice, both *in vivo* and in *vitro*

*KCNT1* epileptic encephalopathy is a childhood genetic disease; therefore, we next tested for evidence of epileptiform activity in pre-adolescent mouse pups by performing acute, head-fixed, unanesthetized *in vivo* electrocorticography on a cohort of *Kcnt1*^m/m^ (n=8) and WT (n=5) pups at postnatal days (P) 13-15. Of the eight *Kcnt1*^m/m^ pups, three showed definite seizure activity (Figure 3A), and all eight exhibited waveforms consistent with IEDs (Figure 3B). Electrographic seizure patterns consisted of evolving patterns of spike and wave discharges that had restricted spatial distribution across the cortical surface (Figure 3A). These seizures lasted from 10 to 280 seconds and stopped spontaneously in all cases (Figure 3C). No WT pups exhibited electrographic seizure activity. Identical offline IED detection algorithms were applied to *Kcnt1*^m/m^ and WT pups, with all *Kcnt1*^m/m^ pups demonstrating a higher occurrence rate of IEDs (Figure 3D). These IEDs were focally distributed over the surface of the cortex across *Kcnt1*^m/m^ pups and, similar to the results from widefield imaging, were more concentrated in anterior regions of the somatomotor cortex (Figure 3E).

Next, we tested whether networks of *Kcnt1*^m/m^ neurons retain their pathological hyperexcitability in an *in vitro* preparation. To do this, we made primary cultures of cortical or hippocampal neurons from *Kcnt1^m/m^* and WT P0 pups and recorded spontaneous spiking activity using multi-electrode array (MEA) analysis. We analyzed a variety of spiking and bursting features between 9 and 27 days *in vitro* (DIV; see Methods). Networks of *Kcnt1^m/m^* and WT neurons matured at a similar rate, as measured by the increase in the number of active electrodes (nAE) per well, which reached a maximum by DIV 17 (not shown), and displayed a variety of activity patterns (Figures 3F). In networks of cortical *Kcnt1^m/m^* neurons, we observed a hyperexcitability phenotype, characterized by an increased mean firing rate (MFR) and an increased bursting frequency (increase in bursts/minute and decrease in interburst interval) (Figure 3G). We also observed an increase in synchrony at the network level, characterized by an increased number of network bursts (Figure 4G). In contrast to cortical networks, there were no significant changes in activity in networks of hippocampal *Kcnt1^m/m^* neurons compared to WT (Figure S3), suggesting regional specificity of neuronal hyperexcitability.

### The *Kcnt1* Y777H variant impairs excitability and AP generation in cortical GABAergic, but not glutamatergic, neurons

The results from the *in vivo* electrophysiology, calcium imaging, and MEA analysis of cortical activity suggest that cortical networks from *Kcnt1^m/m^* mice are hyperexcitable at the end of the second postnatal week, but what are the origins of this hyperexcitability? Previous characterizations in heterologous systems found that the Y796H variant increases peak K_Na_ current from 3- to 11-fold over the WT channel (Tang et al., 2016, Milligan et al., 2014). Because the K_Na_ current has several proposed roles in the regulation of membrane excitability and AP generation, we performed whole-cell current-clamp analysis of cultured cortical neurons during the time frame of network hyperexcitability onset (DIV 13-17). Moreover, because KCNT1 GOF effects are hypothesized to be neuron subtype-specific, we classified the recorded neurons as glutamatergic, fast spiking (FS) GABAergic, or non-fast spiking (NFS) GABAergic, based on expression of GFP driven by the *CaMKII* promoter, AP parameters, and evoked synaptic responses (see Methods).

First, we assessed alterations in the membrane excitability of *Kcnt1^m/m^* glutamatergic neurons. If the K_Na_ current was activated at subthreshold voltages in *Kcnt1^m/m^* neurons, KCNT GOF would be expected to alter the resting membrane potential (V_rest_), input resistance (R_in_), AP threshold, or the minimum amount of current needed to trigger an AP (rheobase); however, none of these parameters were significantly affected by the Y777H variant in glutamatergic neurons (Table S2 and Figures 4A_1_ and 4A_2_). Alternatively, it has been hypothesized that the K_Na_ current activates in response to AP firing, and that KCNT1 GOF increases the rate of AP repolarization in glutamatergic neurons, leading to an increase in AP firing frequency. In this case, KCNT1 GOF would be expected to alter the AP shape; however, the AP half-width, AP repolarization rate, and afterhyperpolarization (AHP) were also not significantly affected by the Y777H variant (Figure 4A_3_ and Table S2). Not surprisingly, the AP firing responses to incremental current injections were also indistinguishable between WT and *Kcnt1^m/m^* glutamatergic neurons (Figure 4A_4_). Thus, despite reported expression of KCNT1 and K_Na_ currents in cortical glutamatergic neurons (Budelli et al., 2009, Rizzi et al., 2016), there were no significant alterations in any of the electrophysiological parameters measured in this neuron type.

An alternative hypothesis is that KCNT1 GOF selectively reduces the excitability of GABAergic neurons. Although, like glutamatergic neurons, the *Kcnt1*^m/m^ FS GABAergic neurons showed no alterations in V_rest_, R_in_, or AP threshold, they did show an increase in the rheobase current (Table S2 and Figures 4B_1_ and 4B_2_), suggesting there may be an increased K_Na_ current near AP threshold levels in this neuron population. Other electrophysiological parameters assessed, including AP half-width, AP repolarization rate, and AHP, were not affected by the Y777H variant in FS GABAergic neurons (Figure 4B_3_ and Table S2). However, incremental current injections showed a decrease in the number of APs per current step in *Kcnt1*^m/m^ FS GABAergic neurons relative to that of WT (Figure 4B_4_).

In contrast to glutamatergic and FS GABAergic neurons, the *Kcnt1*^m/m^ NFS GABAergic neurons showed drastic alterations in membrane properties and AP generation relative to those of WT. The *Kcnt1*^m/m^ NFS GABAergic neurons showed a decrease in R_in_, accompanied by a large increase in the rheobase (Figures 4C_1_ and 4C_2_), whereas the V_rest_, and AP threshold were unchanged (Table S2). The membrane capacitance (C_m_) was also increased in *Kcnt1*^m/m^ NFS neurons (Table S2). There were several other alterations to the AP parameters in *Kcnt1*^m/m^ NFS neurons, including a narrower AP half-width, a faster AP repolarization rate, and a larger AHP (Figure 4C_3_ and Table S2). These data suggest there may be a larger increase in the K_Na_ current in this neuron population, or that the K_Na_ current is increased across a wider voltage range. Importantly, incremental current injections showed a large decrease in the number of APs per current step in *Kcnt1*^m/m^ NFS GABAergic neurons relative to that of WT (Figure 4C_4_). Overall, these data demonstrate that the Y777H variant indeed selectively reduces the intrinsic excitability of FS and, to a greater extent, NFS GABAergic neurons, thus providing a functional answer to the question of how a variant that causes a decrease in excitability could lead to the formation of a hyperexcitable network.

### Layer 2/3 NFS GABAergic neurons in acute cortical *Kcnt1*^m/m^ slices show reductions in membrane excitability and AP generation

Having identified a brain region (Figure 3) and a cell type (Figure 4) that were particularly sensitive to the *Kcnt1* variant, we next wanted to know if the physiological changes observed in culture were present in an experimental preparation with a more organized network. We therefore prepared acute slices containing the anterior motor cortex from WT and *Kcnt1*^m/m^ mice and performed whole-cell recordings from pyramidal neurons (Figure 5A) and GABAergic neurons (Figure 5E) in the medial potion of the slice. Because more superficial cortical layers are enriched for NFS neurons (Tremblay et al., 2016, Lee et al., 2010), and have been reported to show higher levels of KCNT1 expression (Rizzi et al., 2016), we current-clamped neurons in layer 2/3 and injected steps to elicit APs (Figures 5B and 5F). Similar to observations in cortical neuron cultures, *Kcnt1*^m/m^ glutamatergic neurons showed no changes in R_in_ or rheobase relative to those of WT (Figure 5C). They did, however, have a more depolarized AP threshold (Table S3), an effect that is notably opposite of that previously reported from *Kcnt1* knockout mouse models (Martinez-Espinosa et al., 2015, Reijntjes et al., 2019). However, despite their depolarized AP threshold, *Kcnt1*^m/m^ glutamatergic neurons showed similar AP firing patterns to WT glutamatergic neurons (Figure 5D).

**Figure 5.**
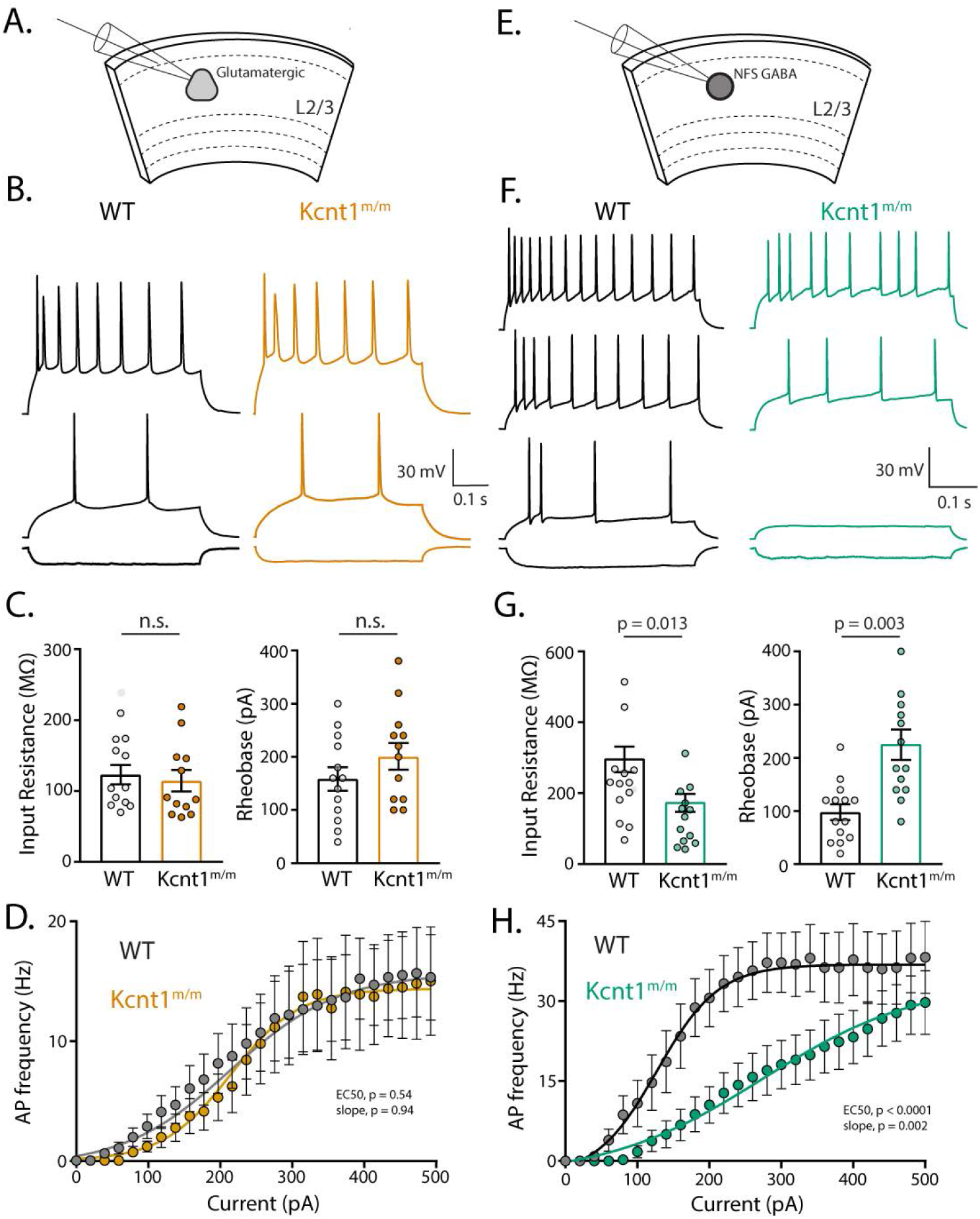
NFS GABAergic neurons in Layer 2/3 of acute slices from motor cortex of *Kcnt1*^m/m^ mice show a strong impairment in AP generation. **(A)** Schematic showing the experimental approach of recording glutamatergic neurons in layer 2/3 of the motor cortex. **(B)** Representative membrane voltage traces of glutamatergic WT (black) and *Kcnt1*^m/m^ (orange) neurons in response to −100, +140, and +240 pA current injections. **(C)** Individual neuron values, mean, and s.e.m. of the input resistance and rheobase of WT and *Kcnt1*^m/m^ glutamatergic neurons. **(D)** Number of APs (mean ± s.e.m.) per current injection step in WT and *Kcnt1*^m/m^ glutamatergic neurons. The EC_50_ (WT: 149 ± 32 pA, *Kcnt1*^m/m^: 180 ± 39 pA) and slope (WT: 117 ± 30 pA, *Kcnt1*^m/m^: 114 ± 32 pA) were not different between genotypes. **(E)** Schematic showing the experimental approach of recording NFS GABAergic neurons in layer 2/3 of the motor cortex. **(F)** Representative membrane voltage traces of NFS GABAergic WT (black) and *Kcnt1*^m/m^ (green) neurons in response to −100, +100, +180, and +300 pA current injections. **(G)** Individual neuron values, mean, and s.e.m. of the input resistance and rheobase of WT and *Kcnt1*^m/m^ NFS GABAergic neurons. **(H)** Number of APs (mean ± s.e.m.) per current injection step in WT and *Kcnt1*^m/m^ NFS GABAergic neurons. The EC_50_ (WT: 136 ± 9 pA, *Kcnt1*^m/m^: 294 ± 20 pA) and slope (WT: 37 ± 9 pA, *Kcnt1*^m/m^: 95 ± 20 pA) were significantly different between genotypes. The lines in the current/AP plots are fits to a Boltzman Sigmoidal curve which were used to determine the EC50 and slope of the relationship. Statistical significance was tested using Generalized Linear Mixed Models.

Importantly, *Kcnt1*^m/m^ NFS GABA neurons showed a significant reduction in R_in_, and large increases in rheobase (Figure 5G) and membrane capacitance (Table S3), all of which were changed in the same manner in cultured neurons, although the AP half-width and AHP were unchanged (Table S3). Moreover, incremental current injections demonstrated significant impairments in AP generation in NFS GABAergic neurons expressing the Y777H variant (Figures 5F and 5H). Thus, the main finding from cultures, that *Kcnt1*^m/m^ NFS GABAergic neurons are most strongly affected by the *Kcnt1* Y777H variant, was also observed in acute slices made from a brain area with demonstrated pathology.

### All *Kcnt1*^m/m^ neurons show an increase in Na^+^-activated K^+^ currents, but the increase in NFS neurons is specific to subthreshold voltages

Why are NFS GABAergic neurons most sensitive to the effects of a variant in a gene that is widely expressed in neurons? To address this question, we recorded Na^+^-activated K^+^ currents (K_Na_) in cultured cortical neurons (Figure 6) by applying voltage step pulses to voltage-clamped neurons and comparing the delayed outward current before and after the addition of TTX, as KCNT1 channels are thought to be the major contributors to this current. Consistent with widespread expression (Saunders et al., 2018, Lein et al., 2007) and previous reports (Budelli et al., 2009), we found significant TTX-sensitive K^+^ currents in cortical glutamatergic neurons that activated at around 0 mV (Figures 6A_1_ and 6A_2_). These currents were also found in WT FS and NFS GABAergic neurons (Figures 6B and 6C), again activating around 0 mV and increasing to peak currents of several nanoamps at +50 mV.

**Figure 6.**
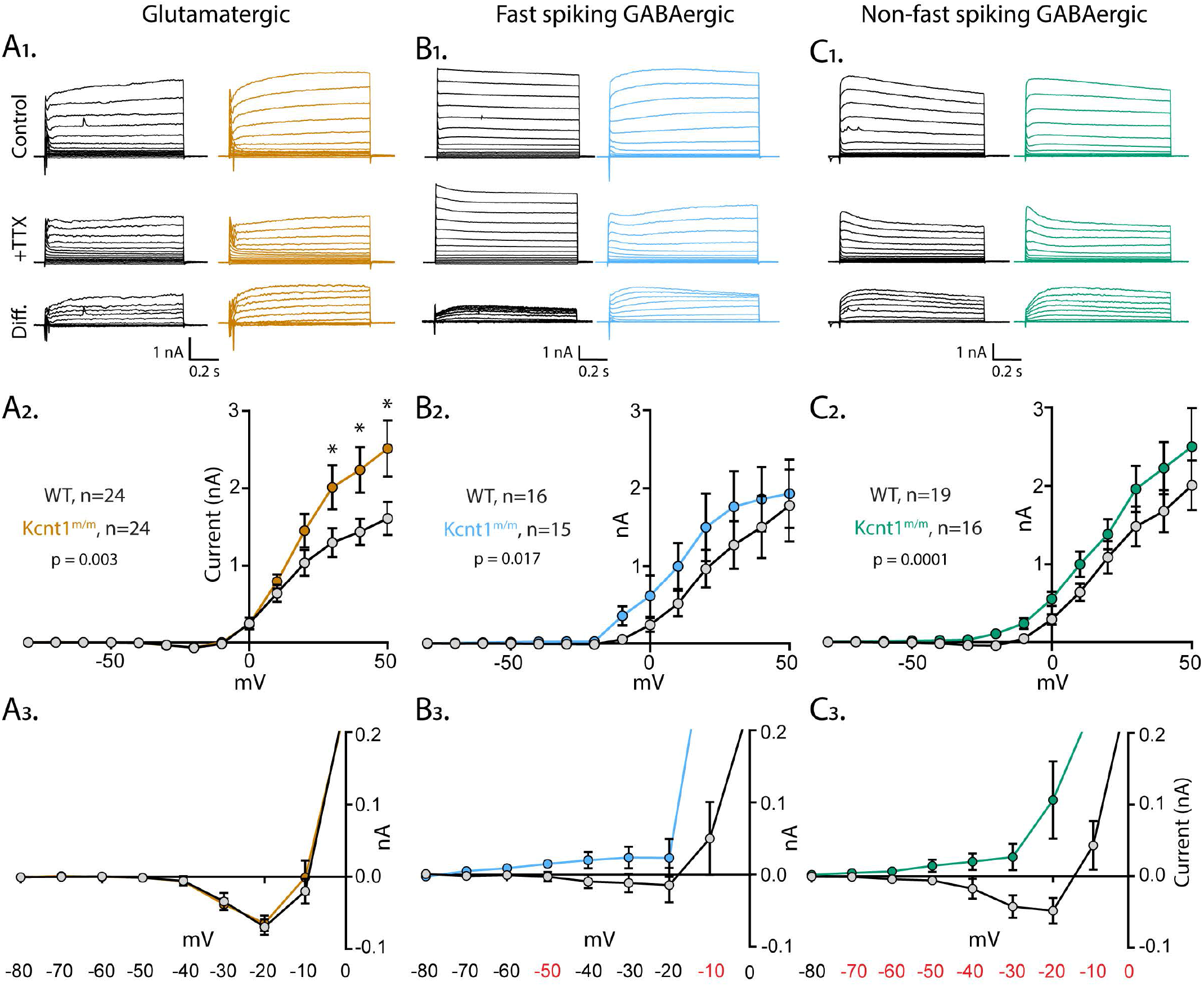
The Y777H variant causes an increase in K_Na_ currents at subthreshold voltages in GABAergic neurons. **(A_1_-C_1_)** Representative traces in control (top), 1 µM TTX (middle), and the difference current (bottom) calculated by subtracting the membrane current response to voltage steps (−80 to +50 mV) from a holding potential of −70 mV in TTX from the response in control external solution in WT (black) and *Kcnt1*^m/m^ (colors) neurons. **(A_2_)** Summary data showing the K_Na_ current (mean ± s.e.m.) for each voltage step in WT (black) and *Kcnt1*^m/m^ (orange) glutamatergic neurons. **(A_3_)** Plot of the K_Na_ current (mean ± s.e.m.) for each voltage step from −80 to 0 mV in WT and *Kcnt1*^m/m^ glutamatergic neurons to illustrate the values that are too small to be seen on the graph in A_2_. **(B_2_)** Summary data showing the K_Na_ current (mean ± s.e.m.) for each voltage step in WT (black) and *Kcnt1*^m/m^ (blue) FS GABAergic neurons. **(B_3_)** Plot of the K_Na_ current (mean and s.e.m.) for each voltage step from −80 to 0 mV in WT and *Kcnt1*^m/m^ FS GABAergic neurons. **(C_2_)** Summary data showing the K_Na_ current (mean ± s.e.m.) for each voltage step in WT (black) and *Kcnt1*^m/m^ (green) NFS GABAergic neurons. **(C_3_)** Plot of the K_Na_ current (mean and s.e.m.) for each voltage step from −80 to 0 mV in WT and *Kcnt1*^m/m^ NFS GABAergic neurons. Statistical significance was tested using Generalized Linear Mixed Models with genotype and current step as fixed effects followed by pairwise comparisons at each level. P values < 0.05 are labeled on the upper plots as asterisks and on the lower plots as red numbers. All other p values were > 0.05.

In all three neuron types, the *Kcnt1* variant significantly increased K_Na_ currents, but the effects were in different voltage ranges in the different cell types. In glutamatergic neurons there was a significant interaction between current step and genotype, indicating an increase in K_Na_ currents at a specific voltage range in *Kcnt1*^m/m^ neurons. Pairwise comparisons at each voltage step detected significant effects (p<0.05) from +30 mV to +50 mV (Figure 6A_2_ and 6A_3_). In FS GABAergic neurons there was also a significant interaction term, but in this case the significant differences between WT and *Kcnt1*^m/m^ neurons were at subthreshold voltages, namely the −50 and −10 mV steps (Figures 6B_2_ and 6B_3_). NFS GABAergic neurons showed an even stronger increase in K_Na_ currents, with a significant interaction term and significant increases at all voltage steps from −70 to 0 mV steps in *Kcnt1*^m/m NFS^ neurons (Figure 6C_3_), demonstrating a large effect across subthreshold voltages. These results are consistent with the observed neuron subtype-specific effects of the KCNT1 GOF variant; more specifically, the large increase in K_Na_ current at subthreshold voltages in *Kcnt1*^m/m^ NFS GABAergic neurons predicts the robust effects observed on reduced membrane excitability, including the decreased R_in_, increased rheobase, and impaired AP firing.

### The *Kcnt1* Y777H variant increases synaptic connections between homotypic neuron pairs

Our finding that KCNT1 GOF caused a selective reduction in excitability in GABAergic neurons is not mutually exclusive with the third proposed hypothesis, which is that *KCNT1* GOF variants cause network excitability because increases in K^+^ current during development alter normal patterns of synaptic connections (Kim and Kaczmarek, 2014). Thus, we tested for altered synaptic connectivity by performing paired recordings of glutamatergic (excitatory, E) and GABAergic (inhibitory, I) neurons and alternatively stimulating each neuron at 0.1 Hz to test baseline connection probability (P_c_) and strength at the four possible motifs (I-I, I-E, E-I, and E-E). P_c_ at the I-E and E-I motifs was not altered in *Kcnt1*^m/m^ networks (Figures 7B and 7C), indicating grossly normal synaptic interactions between glutamatergic and GABAergic neurons. However, P_c_ between both I-I pairs and E-E pairs was higher in *Kcnt1*^m/m^ networks than in those of WT (Figures 7A and 7D). Subclassifying the GABAergic neurons underpowered the analysis, but suggested that both FS and NFS *Kcnt1*^m/m^ neurons show increased inter- and intraconnectivity (data not shown). The amplitudes of the evoked postsynaptic currents (ePSCs) between connected neurons was not significantly different between genotypes for any of the four connection types (Figures 7A-D). Thus, in addition to impairing the excitability of GABAergic neurons, the Y777H variant also increased synaptic connectivity in the network in ways that would be predicted to promote hyperactivity and hypersynchrony.

**Figure 7.**
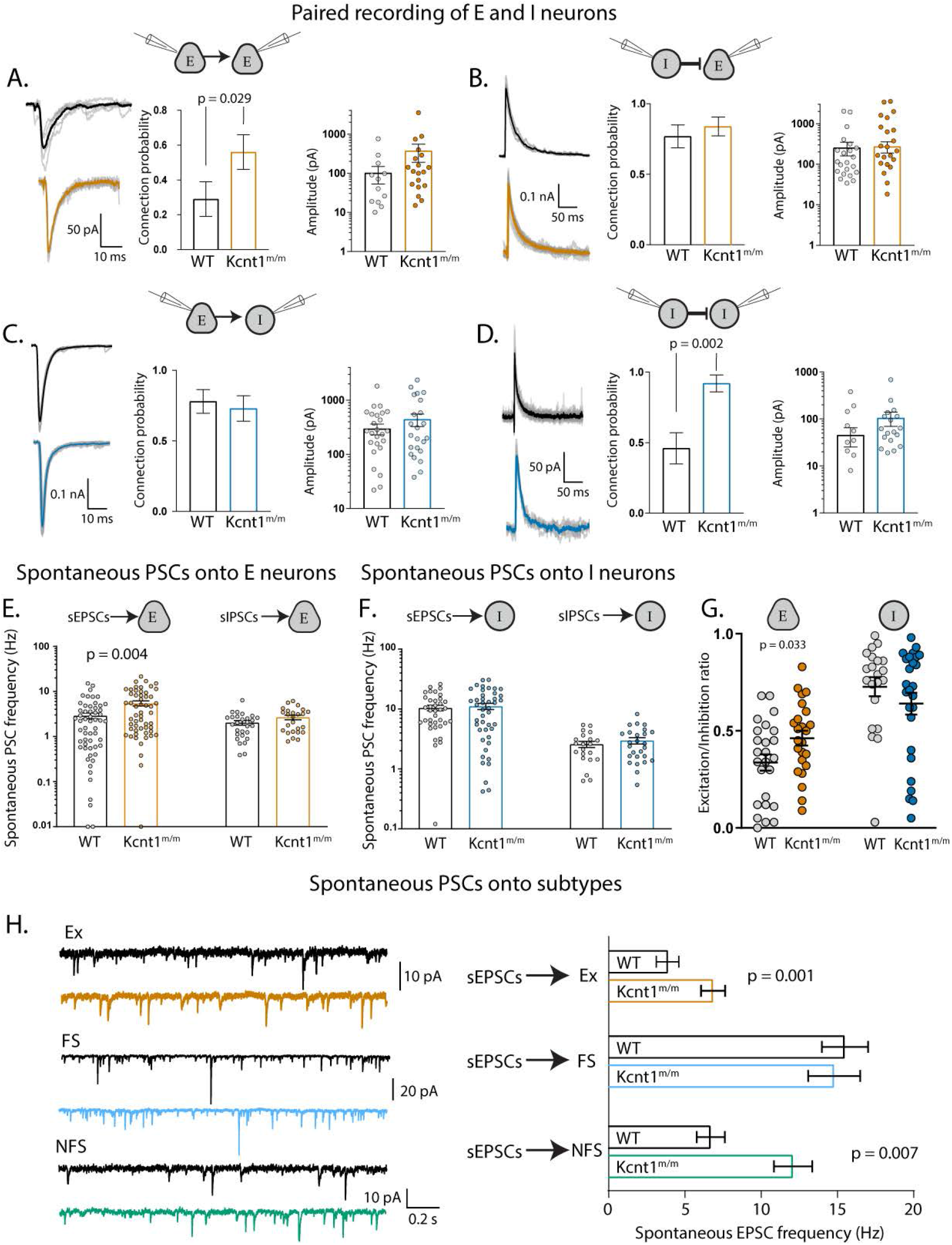
Increases in synaptic connectivity in *Kcnt1*^m/m^ networks contribute to hyperexcitability. **(A_1_-D_1_)** Example traces of synaptic currents evoked by stimulating the neuron type indicated on the left and recording the response in the neuron type indicated on the right. **(A_2_-D_2_)** Summary data (mean ± s.e.m.) of the connection probability between motifs. **(A_3_-D_3_)** Individual pair values and mean ± s.e.m. of peak evoked PSC amplitudes of connected pairs. **(E)** Individual values and mean ± s.e.m. of the spontaneous EPSC (sEPSC) frequency onto glutamatergic neurons (WT, black; *Kcnt1*^m/m^, orange) and GABAergic neurons (WT, black; *Kcnt1*^m/m^, blue). **(F)** Individual values and mean ± s.e.m. of the spontaneous IPSC (sIPSC) frequency onto glutamatergic (WT, black; *Kcnt1*^m/m^, orange) and GABAergic neurons (WT, black; *Kcnt1*^m/m^, blue). **(G)** E/I ratio onto E neurons is increased, but not onto I neurons. **(H_1_)** Example traces of sEPSCs recorded in WT (black) and *Kcnt1*^m/m^(colors) neurons. **(H_2_)** sEPSC frequencies (mean ± s.e.m.) recorded in WT (black) and *Kcnt1*^m/m^ (colors) neurons. Significance values are labeled if p < 0.05 as determined by Generalized Linear Mixed Models.

### The net effect of cellular and synaptic alterations in *Kcnt1*^m/m^ networks is an increase in excitation

Finally, we determined whether the increases in synaptic connectivity were accompanied by alterations in synaptic activity by recording spontaneous PSCs (sEPSCs and sIPSCs) onto voltage-clamped glutamatergic and GABAergic neurons. Similar to the increase in E-E synaptic connectivity, we observed an increase in sEPSC frequency selectively onto *Kcnt1*^m/m^ glutamatergic neurons (Figure 7E). However, sIPSCs were not significantly affected onto *Kcnt1*^m/m^ GABAergic neurons (Figure 7F), possibly due to the offsetting effects of decreased membrane excitability and increased I-I connections. In agreement with the ePSC measurements, the amplitudes and kinetics of the sPSCs were not different between the genotypes onto either neuron group (Table S4). To assess the net effect of altered sPSC activity onto *Kcnt1*^m/m^ neurons, we calculated the E/I ratio, taking into account the relative frequency and size of the sPSCs. The E/I ratio onto *Kcnt1*^m/m^ glutamatergic neurons was significantly higher than that onto WT, whereas the E/I ratio onto GABAergic neurons was similar between the two groups (Figure 7G). Thus, in agreement with the network hyperexcitability, the overall balance of excitation and inhibition in *in vitro Kcnt1*^m/m^ networks is shifted towards excitation.

Although we were initially surprised that the decrease in excitability of *Kcnt1*^m/m^ GABAergic neurons did not lead to decreased sIPSC frequency, it has been shown that for *Scn1a* LOF, reductions in FS (PV-expressing), but not NFS (SST-expressing), neuron excitability lead to decreased sIPSC frequency (Rubinstein et al., 2015). This discrepancy may be due to the fact that a large portion of NFS GABAergic neurons target the distal dendrites of neurons, far from where sIPSCs are measured at the soma. Alternatively, we considered that there might be a compensatory increased excitatory drive specifically onto NFS GABAergic neurons. To test this, we recorded sEPSCs onto E, and FS and NFS GABAergic, neurons (Figure 7H) and found that the sEPSC frequency onto FS neurons was unchanged, whereas the sEPSC frequency onto NFS neurons was increased by the Y777H variant (Figure 7H). This suggests that the decrease in membrane excitability of the *Kcnt1*^m/m^ NFS GABAergic neurons may be partially offset by an increase in excitation onto these neurons, but that this is not the case in FS GABAergic neurons. Thus, although *Kcnt1*^m/m^ glutamatergic and GABAergic neurons and synapses demonstrate complex and likely compensatory alterations, the net effect of these changes is a network with an overall increased excitatory drive.

## DISCUSSION

Classic and recent human genetic studies have provided key insights into the causes of neurodevelopmental disease and severe epilepsies, revealing that genetic variants in ion channels, both ligand- and voltage-gated, comprise approximately one-third of known monogenic causes of seizure disorders (Noebels, 2017). Despite the fact that molecular dysfunction of these channel variants is relatively well-modeled and correctable in heterologous systems, translating these findings to therapies has faced unexpected challenges, likely due to the developmental, cellular, and synaptic complexity of the brain. Progress in tracing the critical steps between ion channel biophysics and pathological synchronization of cortical neurons thus requires precision genetic disease models and multi-level interrogations of their effects (Oyrer et al., 2018, Farrell et al., 2019). To address this, we created a mouse model with a missense mutation in a Na^+^-activated K^+^ channel gene orthologous to a human missense mutation that causes an early-onset seizure disorder (ADNFLE) and has been shown in heterologous systems to be a GOF variant. We showed that homozygous expression of this variant causes an epileptic phenotype with strong parallels to the human disease, and then pursued a systematic interrogation of cellular and network physiology to understand the mechanisms linking channel dysfunction to network abnormalities.

To establish the face validity of our model, and to target our whole-cell electrophysiology experiments to a developmental time window and brain region that we are confident plays a meaningful role in the generation of pathological brain activity, we performed extensive phenotyping of the network pathology using traditional EEG with sleep scoring and *in vivo* widefield Ca^2+^ imaging in adult mice, high-density electrocorticography probes in young mice, and *in vitro* MEA analysis. Using the electrocorticography probes and MEA analysis, we found that *Kcnt1*^m/m^ mice and neurons already showed epileptiform activity by the end of the second postnatal week. Early childhood seizures are a common feature of *KCNT1*-associated, and numerous other, genetic epilepsies. Seizure phenotyping is rarely done at this age in rodents, even though early phenotyping has the potential to uncover seizure phenotypes not present in adults (Lozovaya et al., 2014) and improve our understanding of how well genetic models recapitulate human diseases. Likewise, widefield Ca^2+^ imaging allowed us to pinpoint the M2 region as the cortical source of interictal epileptiform discharges. Our results do not rule out the involvement of other cortical areas in seizure generation, or speak to the strong possibility that cortical interactions with subcortical structures are critical for seizure generation. In fact, the finding that seizures in our model are most likely to happen during NREM sleep, coupled with the fact that ADNFLE is also caused by variants in genes that encode nicotinic acetylcholine receptor subunits (Phillips et al., 1995, Sutor and Zolles, 2001), suggest that thalamic input or cholinergic transmission may play a key role in seizure generation (Klaassen et al., 2006).

By narrowing the pathology to a discrete time and place, we were able to investigate the subtype-specific effects of KCNT1 GOF on neuronal physiology. Results from these studies support a model in which the *Kcnt1* Y777H variant increases the K_Na_ current over different voltage ranges in the three major neuron types of the cerebral cortex; glutamatergic, FS GABAergic, and NFS GABAergic (Figure 6). In GABAergic neurons, because it occurs at subthreshold voltages, the increase is sufficient to impair membrane excitability and AP generation (Figures 4 and 5), changes which are especially strong in the NFS neurons. The reduced excitability of GABAergic neurons is accompanied by increases in E-E and I-I synaptic connectivity, and together, these alterations likely interact to promote both hyperactivity and hypersynchrony in the cortical circuits, especially in the upper layers of cortical region M2, which leads to epileptiform discharges and seizures. This model provides a strong framework for understanding how GOF variants in *KCNT1* lead to epilepsy; however, it remains to be seen if this model will generalize to other ion channels, other GOF variants in K^+^ channels, or even other variants within the *Kcnt1* gene.

Although other K^+^ channel GOF variants have been identified in epilepsy, and neuron subtype-specific effects of these variants have been hypothesized, currently, there are no mouse models to test these hypotheses (Oyrer et al., 2018, Niday and Tzingounis, 2018). For instance, GOF variants have been identified in *KCNMA1* (BK), *KCNA2* (K_V_1.2), *KCNQ2* (K_V_7.2), and *KCNQ3* (K_V_7.3), all of which, similar to KCNT1, show fairly widespread neuronal expression. Moreover, based on their normal functions, they could potentially increase or decrease neuronal excitability by their dual regulation of properties such as resting membrane potential or AP repolarization rate, providing few definitive clues to disease mechanism. Our data provide evidence for the widely held prediction that K^+^ channel GOF causes hyperexcitability by selectively reducing interneuron excitability, a mechanism that may be conserved in other K^+^ channel GOF syndromes, thus demonstrating the effectiveness of using a precision genetic mouse model to probe neuron subtype-specific effects of disease-causing variants.

Prior to this study, only one KCNT1 GOF variant had been characterized in neurons: the MMPSI-associated P942L variant. MMPSI is an earlier onset and more severe form of epilepsy than ADNFLE. Expression of the P942L variant in human induced pluripotent stem cell (iPSC)-derived neurons resulted in an increased K_Na_ current, but only at voltages above +40 mV. Similar to our data, they reported decreased AP width and increased AHP; however, in contrast with our data, the P942L variant caused an increased peak AP firing rate in what are presumably immature glutamatergic neurons (Quraishi et al., 2019). These two studies may have identified different cellular mechanisms because the KCNT1 GOF mechanism is variant- and or disease-dependent. Alternatively, we also observed an increased K_Na_ current at suprathreshold voltages in *Kcnt1*^m/m^ glutamatergic neurons. Thus, it is possible that early effects in cortical glutamatergic neuron excitability have resolved by the time we performed our electrophysiology on more mature glutamatergic neurons, but could contribute to the increase E-E connectivity we observed (Figure 7A).

*Kcnt1* mRNA and protein expression in the CNS are thought to be widespread and present in both glutamatergic and GABAergic neurons, indicating that the differential effects we found in these neuron populations are likely not due to selective expression of KCNT1. In support of this, we found that the Y777H variant increases K_Na_ current in cortical glutamatergic neurons, but that the effects of this increase on physiology are limited because it only occurs at very depolarized voltages. Potential reasons for the lack of a K_Na_ current increase at subthreshold voltages in glutamatergic neurons include glutamatergic-specific alternative splicing of *Kcnt1* mRNA, or coexpression of other channels, such as KCNT2, that can form heteromers with KCNT1 and alter its biophysical properties (Chen et al., 2009, Joiner et al., 1998), thus protecting glutamatergic neurons from the more severe effects of KCNT1 GOF.

Among GABAergic neurons, we identified strong KCNT1 GOF effects on the NFS subtype, which are still electrically, molecularly, and morphologically diverse, with subclasses performing distinct roles in circuit function (Markram et al., 2015). The similar, yet more subtle, effects on the FS subtype, suggesting that there is likely not one particular subclass of inhibitory neurons that is uniquely affected. What cell type differences could account for the stronger effects of KCNT1 GOF on NFS than FS GABAergic neurons? First, FS neurons already show a large K^+^ conductance at subthreshold voltages. This is reflected in their low input resistance, fast time constant, and high rheobase, and caused by high expression of K^+^ leak channels (Okaty et al., 2009). The relatively small effect of KCNT1 GOF on FS neurons may simply reflect the fact that adding a KCNT1-mediated K^+^ conductance on top of an already large K^+^ conductance may have little effect. In support of this, our data show that at 0 mV, the first voltage at which WT neurons show a significant K_Na_ current, the TTX-sensitive current is 10% of the total current in FS neurons, but 20% in NFS neurons.

Second, and perhaps more crucial, is the fact that KCNT1 GOF broadly increases the K_Na_ current throughout subthreshold voltages in NFS neurons, which puts it in a position to have a much stronger effect on AP generation. Although this could have many potential causes, an intriguing hypothesis is that this is due to a larger persistent sodium current (Na_P_) in NFS than FS GABAergic neurons. Strong evidence exists that activation of the K_Na_ current is more heavily dependent on Na_P_ than the transient Na^+^ current (Na_T_), likely due to spatial coupling of the channels, and Na_P_ is active at more hyperpolarized voltages than Na_T_ (Budelli et al., 2009, Hage and Salkoff, 2012). Furthermore, our data show the presence of a TTX-sensitive inward current in WT NFS neurons beginning around −40 mV that is not present in FS neurons (Compare Figure 6B_3_ to 6C_3_), to which Na_P_ is likely a major contributor. Thus, a large Na_P_ in NFS neurons may activate KCNT1 currents at subthreshold voltages, which would cause the dramatic effect of the variant in this cell type.

*KCNT1* and other K^+^ channel GOF epilepsies are typically treatment resistant, and the use of quinidine, which inhibits most *KCNT1* GOF variant currents in heterologous neurons, has shown limited success in treating KCNT1-related epilepsy in humans. Our data show that *Kcnt1*^m/m^ neurons, networks, and mice all showed clear pathology at two weeks of age. In addition to the changes in membrane excitability, which may be correctable by blocking KCNT1 or other ion channel currents, we also found increases in membrane capacitance and synaptic connection probability at multiple synaptic motifs, which likely play a role in the generation of abnormal network activity. These structural changes would be more difficult to reverse by modulating ion channel currents, unless perhaps, the modulation occurred prior to their occurrence, which would argue for earlier treatment with quinidine (Dilena et al., 2018). Alternatively, if the Na_P_ current activates the increased K_Na_ current and results in reduced interneuron excitability, drugs that inhibit Na_P_ could have the dually useful effect of mitigating aberrant activation of the K_Na_ current in NFS GABAergic neurons, while still providing general dampening of membrane excitability in glutamatergic neurons. Na_P_ current inhibitors are currently available and have been used with success in preclinical models of Na^+^ channel dysfunction (Baker et al., 2018, Anderson et al., 2014). Our data suggest that the way forward to designing optimal treatment strategies for genetic epilepsies is a better understanding of the complex interactions between neuron subtype-specific membrane currents, circuit development, and synaptic connections.

## Acknowledgements

This work was supported by NIH/NINDS grants NS087095 (M.C.W.), NS110945 (M.C.W.), NS031348 (W.N.F.), and OD020351 (The Jackson Laboratory Center for Precision Genetics). The authors would like to acknowledge the support of Todd Clason of the Imaging & Physiology Core and the Cellular & Molecular Core at the University of Vermont, and Dr Rod Scott for advice on statistical modeling.

## Author Contributions

Conceptualization, D.B.G., M.J.B., W.N.F., M.C.W.; Methodology, D.W., D.K., M.Y., C.M.L., Formal Analysis, A.N.S., S.C., E.R.C, W.F.T., Y.P., J.N.G., M.C.W.; Investigation, A.N.S., S.C., S.P., E.R.C., W.F.T., S.D., C.D.B., M.A.B., M.C.W.; Writing-Original Draft, A.N.S., S.C., W.F.T., J.N.G., W.N.F., M.C.W.; Writing-Revising and editing A.N.S., S.C., M.J.B., W.N.F., M.C.W.; Supervision, M.Y., J.N.G, D.B.G., M.J.B., W.N.F., M.C.W.; Funding Acquisition, D.B.G., W.N.F., M.C.W.

## MATERIALS AND METHODS

### Mouse husbandry, generation, and genotyping

All mice were bred, and procedures were conducted at the Jackson Laboratory, at Columbia University Irving Medical Center, or at the University of Vermont. Each institution is fully accredited by the Association for Assessment and Accreditation of Laboratory Animal Care, and all protocols were approved by their respective Institutional Animal Care and Use Committees. All experiments were performed in accordance with respective state and federal Animal Welfare Acts and the policies of the Public Health Service. Columbia University protocol numbers: AC-AAAU8484, AC-AAAZ8450, AC-AAAR4414 and AC-AAAU8476 and AC-AAAT6474. University of Vermont protocol numbers: 16-001, 19-034. *Kcnt1*^Y777H^ mice (formal gene and allele symbol: *Kcnt1*^em1(Y777H)Frk^) were generated in the C57BL/6NJ (B6NJ) mouse strain (JAX stock # 005304) using CRISPR/Cas9 and an oligonucleotide donor sequence as part of the Jackson Laboratory Center for Precision Genetics program (JCPG, www.jax.org/research-and-faculty/research-centers/precision-genetics-center), and maintained by backcrossing heterozygous males to wildtype B6NJ females. For some experiments, as noted in the text, male mutant mice were crossed with FVB.129P2-Pde6b+ Tyrc-ch/AntJ dams (JAX stock # 004828) to generate cohorts of N_2_ or F_2_ hybrid mice. Mice were maintained in ventilated cages at controlled temperature (22–23°C), humidity ~60%, and 12h:12h light:dark cycles (lights on at 7:00 AM, off 7:00 PM). Mice had access to regular chow and water, *ad libitum*.

*Kcnt1*^Y777H^ mice were genotyped using PCR amplification primers (*Kcnt1* forward primer: 5’-CTAGGGCTGCAAACACAACA-3’; *Kcnt1* reverse primer: 5’-TCAAGCAGCAACACGATAGG-3’) with standard thermocycler amplification conditions, and the annealing temperature set at 58°C. Following amplification, a restriction cut was performed with the enzyme *Nla*III to distinguish mutant (127 and 44 bp products after cut) from wild-type alleles (171 bp product).

### Western Blot

For crude membrane/cytosol fractioning, cortices were homogenized in the following homogenization buffer: 0.32 M sucrose, 10 mM HEPES pH 7.4, 2mM EDTA in H2O, containing a protease inhibitor cocktail and a phosphatase inhibitor cocktail. The samples were homogenized with a motor-driven homogenizer and centrifuged at 1000 × g for 15 min at 4°C to remove the pelleted nuclear fraction. The supernatant was further centrifuged at 16000 rpm for 20 min at 4°C to yield the cytosolic fraction in the supernatant and the pelleted membrane fraction. The pellet was resuspended in homogenization buffer. Protein concentrations were determined with the Pierce BCA Protein Assay Kit (Thermo Fisher Scientific #23225) using BSA as a standard.

Sample protein (10 μg) was mixed with 4X LDS Sample Buffer (NuPAGE), and 10X Sample Reducing Agent (NuPAGE), heated for 10 min at 70°C, separated on 4-12% Bis-Tris Mini Gel (NuPAGE) and transferred to a PVDF membrane (Millipore #ISEQ00010). Nonspecific binding was blocked for 1 hr at RT with 5% BLOTTO (Bio-Rad) in Tris-buffered saline with 0.1% Tween (TBST). Membranes were incubated overnight at 4°C with the primary antibody mouse anti-KCNT1 (1:1000, Abcam #ab94578), washed 3 × 10 min with TBST and incubated with the secondary antibody HRP-conjugated goat anti-mouse (1:10000, Santa Cruz Biotechnology #sc-2005) in 5% BLOTTO for 1 hr at RT and washed again 3 × 10 min with TBST. Blots were incubated for 5 min with Pierce ECL Western Blotting Substrate (Thermo Fisher Scientific #32106) and developed with a Kodak X-OMAT 2000A Processor. Blots were stripped for 15 min at RT in ReBlot Plus Strong Solution (Millipore #2504), blocked for 30 min in 5% BLOTTO, incubated for 1 hr at RT with an HRP-conjugated mouse anti-β-actin antibody (1:5000, Santa Cruz Biotechnology #sc-47778) in 5% BLOTTO and developed as previously described. β-actin was used as a loading control.

### Behavior

#### Open field exploration

Each mouse was gently placed in the center of a clear Plexiglas arena (7.31 x 27.31 x 20.32 cm, Med Associates ENV-510) lit with dim light (~5 lux), and allowed to ambulate freely for 60 min. Infrared (IR) beams embedded along the X, Y, Z axes of the arena automatically track distance moved, horizontal movement, vertical movement, stereotypies, and time spent in center zone. At the end of the test, the mouse was returned to the home cage and the arena was cleaned with 70% ethanol followed by water drying.

#### Elevated plus maze

The elevated plus maze test was conducted as described previously (Yang et al., 2012). The elevated plus-maze consisted of two open arms (30cm x 5cm) and two closed arms (30 x 5 x 15 cm) extending from a central area (5 x 5 cm). Photo beams embedded at arm entrances registered movements. Room illumination was approximately 5 lux. The test began by placing the subject mouse in the center, facing a closed arm. The mouse was allowed to freely explore the maze for 5 min. Time spent in the open arms and closed arms, the junction, and number of entries into the open arms and closed arms, were automatically scored by the MED-PC V 64bit Software (Med Associates). At the end of the test, the mouse was gently removed from the maze and returned to its home cage. The maze was cleaned with 70% ethanol and wiped dry between subjects.

#### Acoustic startle response

Acoustic startle response was tested using the SR-Laboratory System (San Diego Instruments, San Diego, CA) as described previously (Yang et al., 2012). Test sessions began by placing the mouse in the Plexiglas holding cylinder for a 5-min acclimation period. For the next 8 min, mice were presented with each of six trial types across six discrete blocks of trials, for a total of 36 trials. The intertrial interval was 10–20 s. One trial type measured the response to no stimulus (baseline movement). The other five trial types measured startle responses to 40 ms sound bursts of 80, 90, 100, 110 or 120 dB. The six trial types were presented in pseudorandom order such that each trial type was presented once within a block of six trials. Startle amplitude was measured every 1 ms over a 65 ms period beginning at the onset of the startle stimulus. The maximum startle amplitude over this sampling period was taken as the dependent variable. Background noise level of 70 dB was maintained over the duration of the test session.

#### Fear Conditioning

Training and conditioning tests were conducted in two identical chambers (Med Associates) that were calibrated to deliver identical footshocks. Each chamber was 30 cm × 24 cm × 21 cm with a clear polycarbonate front wall, two stainless side walls, and a white opaque back wall. The bottom of the chamber consisted of a removable grid floor with a waste pan underneath. When placed in the chamber, the grid floor connected with a circuit board for delivery of scrambled electric shock. Each conditioning chamber was inside a sound-attenuating environmental chamber (Med Associates). A camera mounted on the front door of the environmental chamber recorded test sessions, which were later scored automatically, using the VideoFreeze software (Med Associates). For the training session, each chamber was illuminated with a white house light. An olfactory cue was added by dabbing a drop of imitation almond flavoring solution (1:100 dilution in water) on the metal tray beneath the grid floor. The mouse was placed in the test chamber and allowed to explore freely for 2 min. A pure tone (5kHz, 80 dB), which serves as the conditioned stimulus (CS), was played for 30 s. During the last 2 s of the tone, a footshock (0.5 mA) was delivered as the unconditioned stimulus (US). Each mouse received three CS-US pairings, separated by 90 s intervals. After the last CS-US pairing, the mouse remained in the chamber for another 120 s, during which freezing behavior was scored by the VideoFreeze software. The mouse was then returned to its home cage. Contextual conditioning was tested 24 h later in the same chamber, with the same illumination and olfactory cue present but without footshock. Each mouse was placed in the chamber for 5 min, in the absence of CS and US, during which freezing was scored, and then returned to its home cage. Cued conditioning was conducted 48 h after training. Contextual cues were altered by covering the grid floor with a smooth white plastic sheet, inserting a piece of black plastic sheet bent to form a vaulted ceiling, using near infrared light instead of white light, and dabbing vanilla instead of banana odor on the floor. The session consisted of a 3 min free exploration period followed by 3 min of the identical CS tone (5kHz, 80dB). Freezing was scored during both 3 min segments. The mouse was then returned to its home cage and the chamber thoroughly cleaned of odors between sessions, using 70% ethanol and water.

#### Nesting

The nesting test was performed as previously described (Deacon, 2006). Briefly, mice were placed in individual cages at 6 pm, with a piece of nestlet weighing 2.5 g. The next morning, the remaining unshredded nestlet was weighed and a nesting score between 1 (no nest) and 5 (perfect nest) was attributed depending on nest quality.

### Video-EEG

Electrode implantation of adult mice was performed surgically essentially as recently described (Asinof et al., 2016). Mice were anesthetized with tribromoethanol (250 mg/kg i.p., Sigma-Aldrich #T48402). Three small burr holes were drilled in the skull (1 mm rostral to the bregma on both sides and 2 mm caudal to the bregma on the left) 2 mm lateral to the midline. One hole was drilled over the cerebellum as a reference. Using four teflon-coated silver wires soldered onto the pins of a microconnector (Mouser electronics #575-501101) the wires were placed between the dura and the brain and a dental cap was then applied. The mice were given a post-operative analgesic of carprofen (5 mg/kg subcutaneous Rimadyl injectable) and allowed a 48-hr recovery period before recordings were taken. To record EEG signals, mice were connected to commutators (PlasticOne) with flexible recording cables to allow unrestricted movements within the cage. Signal (200 samples/s) was acquired on a Grael II 48 EEG amplifier (Compumedics), and the data was examined in Profusion 5 software (Compumedics). Differential amplification recordings were recorded pair-wise between all three electrodes, as well as referential, providing a montage of 6 channels for each mouse. Mouse activity was captured simultaneously by video monitoring using a Sony IPELA EP550 model camera, with an infrared light to allow recordings in the dark. We recorded continuously for 24 to 72 hours.

For sleep analysis, we used a custom-written Matlab program to classify wake, NREM sleep, REM sleep and detect GTCS and TS events. The program first used fast Fourier transform (FFT) to calculate the power spectrum of the EEG using a 5-sec sliding window, sequentially shifted by 2-sec increments. Then it semi-automatedly classified into different brain states using the following criteria: NREM sleep, high power at low-frequency (1-4 Hz) and low EMG activity; REM sleep, high power at theta frequencies (6-9 Hz) and low EMG activity; GTCS and TS, high power at “seizure”-frequencies (19-23 Hz); wake, the remaining (or default) state. We chose the 19-23 Hz band to detect seizures based on its clear separation from normal brain oscillatory activities. The classification was followed by manual inspection to further refine the scoring. Brain states prior to each GTCS/TS event were used in Table S1.

### Widefield Calcium Imaging

Mice that were heterozygous for the *Snap25*-GCaMP6s construct (Madisen et al., 2015) (Jackson Labs Stock No: 025111) and either homozygous Y777H or wildtype at the *Kcnt1* locus were generated. To gain optical access to the dorsal cortex, the animal’s scalp and the underlying periosteum were removed before coating the dorsal skull with UV cure cyanoacrylate (Loctite 4305) and attaching a custom aluminum headplate to the skull. This headplate framed the glue covered skull and allowed the animal to be secured on a 20 cm diameter Styrofoam treadwheel during imaging sessions. In one homozygous mutant mouse the dorsal skull was removed entirely and replaced with a glass window (Labmaker, Crystal Skull) following a previously described protocol (Kim et al., 2016). Imaging was performed using a custom tandem-lens epifluorescence macroscope built according to a design previously described (Wekselblatt et al., 2016). This macroscope was configured for 2.1X magnification using a Nikon Nikkor 50mm f/1.2 lens paired with a Nikon 105mm f/1.8 lens. During all sessions, images were acquired at 40Hz with an Andor Zyla 5.5 10-tap sCMOS camera set to 4 x 4 pixel binning. Successive frames were illuminated with blue and green light and the green reflectance images were used to estimate changes in fluorescence resulting from hemodynamic factors and correct for them in GCaMP images. For blue illumination, we used an X-Cite 120 LED (Lumen Dynamics, XT120L) with a filter set consisting of an ET470/40x excitation filter, a T495lpxr dichroic, and an ET525/50m emission filter (Set 49002, Chroma Technology, Bellows Falls, VT). Total blue light power delivered to the implant typically ranged from 18-21 mW. For green illumination we used a 530nm 6.8mW Thorlabs LED (M530F2) driven by a Thorlabs tcube (LEDD1B) coupled to a fiber optic obliquely pointed at the brain. To switch illumination sources between frames we relied on an Arduino Uno R3 (A000066) microcontroller operating on the Zyla ‘expose’ signal. At the beginning of each session we exposed a dark frame in which no illumination was used. This frame was subtracted from all illuminated frames in the session as the first processing step. Subsequently, reflectance and GCaMP images were deinterleaved and linearly interpolated from 20 to 40 Hz. We calculated ΔF/F for each pixel in the image that covered the brain, as defined by a hand-drawn mask. We estimated F for each pixel as the average of the bottom 20th percentile of fluorescent values over the entire session. Following ΔF/F calculation we subtracted fractional green fluorescence (ΔF/F_mean_) with F_mean_ calculated over the full session. Finally, we performed Singlular Value Decomposition (SVD) on the ΔF/F image stacks and reconstructed images using the first 50 singular values for all subsequent analysis. The SVD code was a modification of that provided by the Cortex Lab group at UCL (github address: https://github.com/cortex-lab/widefield). To rigidly align area borders derived from the Allen Common Coordinate Framework v3 to each brain we relied on the landmark based strategy previously described (Musall et al., 2019).

### Mouse pup EEG

Eight *Kcnt1*^m/m^and five WT pups at postnatal days 13-15 were used for in vivo electrophysiology. All experiments were performed in accordance with protocols approved by IACUC at Columbia University Irving Medical Center. Pups were anesthetized using isoflurane and electromyography (EMG) electrodes were placed for real-time monitoring of respiratory rate, heart rate, and muscle activity. The animals were then given systemic and local analgesia, head-fixed, and a craniotomy was made over the dorsal cortical surface of one hemisphere. This craniotomy was centered over somatomotor cortex, spanning 2.5 mm in the mediolateral direction and 3.5 mm in the anterioposterior direction between bregma and lambda. A NeuroGrid (ultra-conformable, biocompatible surface electrocorticography array, 119 electrodes, 177 µm pitch) array was placed on the surface of the dura and covered with a piece of sterile compressed sponge. The pup was transferred to a temperature and humidity-controlled enclosure and allowed to recover for 30 minutes, after which recording commenced. Signals were amplified, digitized continuously at 20 kHz using a head-stage directly attached to the NeuroGrid (RHD2000, Intan technology), and stored for off-line analysis with 16-bit format. Data were analyzed using MATLAB (MathWorks) and visualized using Neuroscope. At the completion of electrophysiological recording, pups were euthanized.

Neurophysiologic recordings were synchronized with the EMG signals to facilitate identification and elimination of any epochs with artifacts from subsequent analysis. Ictal patterns were identified using a line length algorithm and confirmed with visual screening of the raw data. Interictal epileptiform discharges were detected using previously employed frequency and duration features (Gelinas et al., 2016).

### Multi-electrode arrays (MEA)

#### Primary neuron culture

One to seven days before dissection, 48-well MEA plates (Axion Biosystems #M768-KAP-48) were coated with 50 µg/mL poly-D-lysine (Sigma-Aldrich #P0899-50MG) in borate buffer, then washed three times with phosphate buffered saline (PBS) and stored in PBS at 4°C until use. Prior to use, PBS was aspirated and plates were allowed to dry in a sterilized hood. Cortical or hippocampal neurons were dissociated from the brains of postnatal day 0 (P0) C57BL/6NJ WT or *Kcnt1*^m/m^mice. Pups were decapitated, weighed and genotyped. The entire cerebral cortex was rapidly dissected and cut into small pieces under sterile conditions in cold Hibernate A solution (Gibco, #A1247501). Cortices from two WT or *Kcnt1*^m/m^pups were pooled together. The dissected cortices were then enzymatically digested in 20 U mL^−1^ Papain plus DNAse (Worthington Biochemical Corporation, #LK003178 and #LK003172) diluted in Hibernate A for 20 min at 37°C. Cells were pelleted by centrifugation at 300RCF for 5 min, then the digestion was neutralized by aspirating off the supernatant and adding warm Hibernate A media. Cells were mechanically dissociated by trituration, and counted using a hemocytometer with Trypan blue counterstain. Cells were pelleted by centrifugation at 300RCF for 5 min and re-suspended at a density of 6,000 cells/µl in warm Neurobasal-A (Gibco #10888022) + 1X B27 supplement (Gibco #17504044) + 1X GlutaMax (Gibco #35050061) + 1% HEPES (Gibco #15630080) + 1% Penicillin/Streptinomycin (Gibco # 15140122) + 1% fetal bovine serum (Gibco #26140079) + 5 ug/mL Laminin (Sigma-Aldrich #L2020). 50,000 cells were plated on a pre-coated 48-well MEA plate in a 40 uL drop. The day after plating (DIV1), 100% of the media was removed and replaced with warm Neurobasal-A + 1X B27 supplement + 1X GlutaMax + 1% HEPES + 1% Penicillin/Streptinomycin (NBA/B27 medium). Glial growth was not chemically suppressed. Cultures were maintained at 37°C in 5% CO_2_. Media was 50% changed every other day with fresh warm NBA/B27 starting on DIV3, after each recording session.

#### Data analysis of spontaneous recordings

MEA recordings were conducted on media change days prior to media change starting on DIV5. Plates were equilibrated for 5 minutes then recorded for 15 minutes per day using Axion Biosystems Maestro 768 channel amplifier at 37°C in a CO_2_ gas-controlled chamber and Axion Integrated Studios (AxIS) software v2.4. Each well on a 48-well plate is comprised of 16 electrodes on a 4 by 4 grid with each electrode capturing activity of nearby neurons. A Butterworth band-pass filter (200-3000Hz) and an adaptive threshold spike detector set at 7x the standard deviation of the noise was used during recordings. Raw data and a spike list files were collected. Spike list files were used to extract additional spike, burst, and network features, using a custom MEA analysis software package for interpretation of neuronal activity patterns, meaRtools, based on rigorous permutation statistics that enables enhanced identification of over 70 activity features (Gelfman et al., 2018). Specifically, we analyzed spiking and bursting rates, burst duration, and the time between bursts (i.e. interburst interval, IBI), as well as synchronicity of the network. We determined the parameters for detecting neuronal bursts and network events based on published reports and experimentation (Mack et al., 2014, McConnell et al., 2012). Activity data was inspected to remove inactive electrodes and wells. In order for an electrode to be considered active, we required that at least five spikes per minute were recorded. Wells in which fewer than 4 electrodes were active for > 50% of the days of recording were considered inactive and removed from analyses. For synchronous network events, at least 5 electrodes (> 25% of the total in a well) were required to participate in a network event in order for the network event to qualify as a network spike or burst. Events with less participating electrodes were filtered. Bursts were detected using the Maximum Interval burst detection algorithm (Neuroexplorer software, Nex Technologies) implemented in the meaRtools package (Gelfman et al., 2018). We required that a burst consists of at least 5 spikes and lasts at least 0.05 second, and that the maximum duration between two spikes within a burst be 0.1 second at the beginning of a burst and 0.25 second at the end of a burst. Adjacent bursts were further merged if the duration between them is less than 0.8 second. These parameters were chosen based on the literature and on in-house experimentation (Mack et al., 2014). To analyze data over time, we performed permuted Mann-Whitney U tests. The values for each well for the chosen DIVs were combined and a Mann-Whitney U (MWU) test was performed. The labels for each well (WT vs *Kcnt1*^m/m^) were then shuffled and permuted 1,000 times to create 1,000 data sets that were tested for significance using a MWU test. We report the permuted p-values as the rank of the true p-value within the distribution of permuted p-values. We also report the combined p-value of the plates, calculated using an R script developed in-house.

### Whole cell electrophysiology

#### Cell culture

Conventional primary cultures were grown on astrocytes derived from wild-type C57BL/6J mice (Jackson Labs stock: 000664), as previously described (Barrows et al., 2017). Cortices were dissected from postnatal day 0-1 (P0-P1) mice of either sex and placed in 0.05% trypsin-EDTA (Gibco) for 15 min at 37°C in a Thermomixer (Eppendorf) with gentle agitation (800 rpm). Then, the cortices were mechanically dissociated with a 1 mL pipette tip and the cells were plated into T-75 flasks containing astrocyte media [DMEM media supplemented with glutamine (Gibco) and MITO+ Serum Extender (Corning). After the astrocytes reached confluency, they were washed with PBS (Gibco) and incubated for 5 min in 0.05% trypsin-EDTA at 37°C, and then resuspended in astrocyte media. Astrocytes were added to 6-well plates containing 25-mm coverslips precoated with coating mixture [0.7 mg/ml collagen I (Corning) and 0.1 mg/ml poly-D-lysine (Sigma) in 10 mM acetic acid].

For the primary neuron culture, the dorsomedial cortices from P0-P1 mice of both sexes were dissected in cold HBSS (Gibco). The tissue was then digested with papain (Worthington) for 60-75 min and treated with inactivating solution (Worthington) for 10 min, both while shaking at 800 rpm at 37°C in a Thermomixer. The neurons were then mechanically dissociated and counted. 150,000 neurons/well were added to 6-well plates in NBA plus [Neurobasal-A medium (Gibco) supplemented with Glutamax (Gibco) and B27 (Invitrogen)]. After plating, approximately 4 × 10^10^ genome copies (GC) of AAV8-CaMKII-GFP (UNC Vector Core) was added to each well.

#### Culture Electrophysiology

Whole-cell recordings were performed with patch-clamp amplifiers (MultiClamp 700B; Molecular Devices) under the control of Clampex 10.3 or 10.5 (Molecular Devices, pClamp, RRID:SCR_011323). Data were acquired at 20 kHz and low-pass filtered at 6 kHz. The series resistance was compensated at 70%, and only cells with series resistances maintained at less than 15 MΩ were analyzed. Patch electrodes were pulled from 1.5-mm o.d. thin-walled glass capillaries (Sutter Instruments in five stages on a micropipette puller (model P-97; Sutter Instruments). Internal solution contained the following: 136 mM K-gluconate, 17.8 mM HEPES, 1 mM EGTA, 0.6 mM MgCl_2_, 4 mM ATP, 0.3 mM GTP, 12 mM creatine phosphate, and 50 U/ml phosphocreatine kinase or 136 mm KCl, 17.8 mm HEPES, 1 mm EGTA, 0.6 mm MgCl2, 4 mm ATP, 0.3 mm GTP, 12 mm creatine phosphate, and 50 U/ml phosphocreatine kinase. The pipette resistance was between 2 and 4 MΩ. Standard extracellular solution contained the following (in mM): 140 NaCl, 2.4 KCl, 10 HEPES, 10 glucose, 4 MgCl_2_, and 2 CaCl_2_ (pH 7.3, 305 mOsm). All experiments were performed at room temperature (22–23°C). Whole-cell recordings were performed on neurons from control and mutant groups in parallel on the same day (day 13–16 *in vitro*). All experiments were performed by two independent investigators blinded to the genotypes.

For current-clamp experiments, intrinsic electrophysiological properties of neurons were tested by injecting 500-ms square current pulses incrementing in 20 pA steps, starting at −100 pA. Resting membrane potential (V_m_) was calculated from a 50 ms average before current injection. The membrane time constant (τ) was calculated from an exponential fit of current stimulus offset. Input resistance was calculated from the steady state of the voltage responses to the hyperpolarizing current steps. Membrane capacitance was calculated by dividing the time constant by the input resistance. Action potentials (APs) were evoked with 0.5 s, 20 pA depolarizing current steps. AP threshold was defined as the membrane potential at the inflection point of the rising phase of the AP. AP amplitude was defined as the difference in membrane potential between the AP peak and threshold and the afterhyperpolarization was the difference between AP threshold and the lowest V_m_ value within 50 ms. The AP half-width was defined as the width of the AP at half-maximal amplitude. The maximum depolarization and repolarization rates were determined by differentiating the AP waveform and finding the peak values. To obtain the neuron’s maximum firing frequency, depolarizing currents in 20-pA steps were injected until the number of APs per stimulus reached a plateau phase. Rheobase was defined as the minimum current required to evoke an AP during the 500 ms of sustained somatic current injections. AP half-width adaptation was determined by dividing the half-width of the last AP of a train at steady state frequency by the first. AP frequency adaptation was determined by dividing the mean interspike interval of the last four APs by the minimum. The membrane potential values were not corrected for the liquid junction potential. GABAergic neurons were classified as fast spiking (FS) if their maximum mean firing rate reached above 60 Hz and their AP half-widths increased by less than 25% during (Casale et al., 2015, Avermann et al., 2012). All others were considered non-fast spiking (NFS).

For voltage-clamp experiments to measure the sodium-activated K^+^ current, neurons were held at −70 mV and given 1 s voltage pulses in 10 mV steps over a range of −80 to +50 mV. Recordings were obtained for each cell in standard extracellular solution or extracellular solution containing 0.5 µM Tetrodotoxin (TTX). Current traces from the TTX solution were subtracted from the current traces obtained from the standard solution. The difference current over the 100 ms at the end of the voltage pulse was considered the steady state K_Na_ current.

For voltage-clamp experiments to measure synaptic currents, neurons were held at −70 mV, except for evoked IPSC measurements, for which neurons were held at −30 mV. AP-evoked EPSCs were triggered by a 2 ms somatic depolarization to 0 mV. The shape of the evoked response, the reversal potential and the effect of receptor antagonists [10 μM NBQX (Tocris Bioscience) or 20 μM bicuculline (BIC, Tocris Bioscience)] were analyzed to verify the glutamatergic or GABAergic identities of the currents. The E/I ratio was calculated as the product of the sEPSC frequency and charge over the sum of the sEPSC frequency and charge and the product of the sIPSC frequency and charge. Neurons were stimulated at 0.1 Hz in standard external solution to measure basal-evoked synaptic responses. Electrophysiology data were analyzed offline with AxoGraph X software (AxoGraph Scientific, RRID:SCR_014284).

#### Slice Electrophysiology

P20–P29 mice were deeply anesthetized with isoflurane and decapitated. The brain was removed and 350 µm coronal slices of frontal cerebral cortex were cut in ice-cold cutting solution (126 mM NaCl, 25 mM NaHCO_3_, 10 mM d-glucose, 3.5 mM KCl, 1.5 mM NaH_2_PO_4_, 0.5 mM CaCl_2_, 10.0 mM MgCl_2_) with a Leica VT1000S. Slices were then transferred to a storage chamber with fresh artificial cerebrospinal fluid (aCSF) containing 126 mM NaCl, 3.5 mM KCl, 1.0 mM MgCl_2_, 2.0 mM CaCl_2_, 1.5 mM NaH_2_PO_4_, 25 mM NaHCO_3_, and 10 mM d-glucose, pH 7.3-7.4, and were incubated at 37 °C for 30 min. The slices then were incubated at room temperature for at least another 30 min before recording. All solutions were continuously bubbled with 95% O_2_ and 5% CO_2_.

Whole-cell current-clamp and voltage-clamp recordings were obtained from cortical layer 2/3 pyramidal cells and interneurons at 32°C using a Multiclamp 700B and Clampex 10.5 software. Individual slices were transferred to a recording chamber located on an upright microscope (BX51; Olympus) and were perfused with oxygenated aCSF (2 mL/min). Neurons were visualized using IR-differential interference contrast microscopy. Layer 2/3 pyramidal cells were distinguished from interneurons by their triangular morphology, large soma, and pronounced apical dendrite. NFS GABAergic cells were distinguished from pyramidal neurons by their typical ovoid-shaped cell bodies and were finally confirmed by electrophysiological recordings as displaying shorter AP half widths and higher peak AP frequency (Avermann et al., 2012). Intracellular solution contained (in mM): 136 mM K-gluconate, 17.8 mM HEPES, 1 mM EGTA, 0.6 mM MgCl2, 4 mM ATP, 0.3 mM GTP, 12 mM creatine phosphate, and 50 U/ml phosphocreatine kinase, pH 7.2. When patch electrodes were filled with intracellular solution, their resistance ranged from 4–6 MΩ. Access resistance was monitored continuously for each cell. The protocols and determination of electrophysiological parameters were conducted as described above for the culture recordings. All experiments were performed and analyzed by an investigator blinded to the animal genotypes.

### Experimental Design and Statistics

Prism 7 (GraphPad Prism, RRID:SCR_002798) was used to perform statistical tests on adult behavioral tests and to create all graphs. Multiple group comparisons were done using two-tailed t-test with correction using the Holm-Sidak method, Mann-Whitney U test, or two-way repeated measures ANOVA with correction using the Sidak’s test, as indicated. Statistics for MEA data and seizure frequency were performed in R using a Mann-Whitney U test with 1,000 permutations.

To test for statistical significance for all whole cell electrophysiology experiments, we used generalized linear mixed models (GLMM) in SPSS (26.0 Chicago, III (IBM, RRID:SCR_002865), which allows for within-subject correlations and the specification of the most appropriate distribution for the data. Because neurons and animals from the same culture or animal are not independent measurements, culture or litter was used as the subject variable, and animals and neurons were considered within-subject measurements. All data distributions were assessed with the Shapiro-Wilk test. Datasets that were significantly different from the normal distribution (p < 0.05) were fit with models using the gamma distribution and a log link function, except for synaptic connection probability, which was fit with the binomial distribution and probit link. Normal datasets were fit with models using a linear distribution and identity link. We used the model-based estimator for the covariance matrix and goodness of fit was determined using the corrected quasi likelihood under independence model criterion and by the visual assessment of residuals. All values reported in the text, figures, and tables are estimated marginal means +/− standard error.

---

**Table S1.** Seizures in Kcnt1 mutant mice predominately happen during NREM sleep.

| Mouse ID | Number of GTCS or TS |  |  |  | sleep analysis |  |  |
| --- | --- | --- | --- | --- | --- | --- | --- |
|  | Total | wake | NREM | REM | wake<br>(min/24h) | NREM<br>(min/24h) | REM<br>(min/24h) |
| 1 | 25 | 0 | 23 | 2 | 677.8 | 693.3 | 49.65 |
| 2 | 0 | 0 | 0 | 0 | 711.3 | 666.5 | 50.3 |
| 3 | 11 | 0 | 9 | 2 | 776.1 | 598.6 | 64.8 |
| 4 | 19 | 0 | 19 | 0 | 798.9 | 601.1 | 30.5 |
| 5 | 12 | 2 | 7 | 3 | 681.9 | 671.6 | 86.6 |
| 6 | 46 | 1 | 45 | 0 | 655.55 | 747.55 | 27.55 |
| 7 | 32 | 3 | 28 | 1 | 616 | 765.65 | 45.95 |

Shown are the numbers of GTCS or TS events in wake, NREM sleep, and REM sleep in 7 Kcnt1 homozygous mutant mice. Note the majority of seizure events occurred during NREM sleep. EEG and EMG were recorded for 1-3 days. The amount of wake, NREM sleep, and REM sleep time were normalized to a 24-hour period.

**Table S2.** Electrophysiological parameters of current clamp recordings from neuronal cultures.

|  | Glutamatergic Neurons |  |  | Fast Spiking GABA |  |  | non-Fast Spiking GABA |  |  |
| --- | --- | --- | --- | --- | --- | --- | --- | --- | --- |
|  | WT | Kcnt1<sup>m/m</sup> |  | WT | Kcnt1<sup>m/m</sup> |  | WT | Kcnt1<sup>m/m</sup> |  |
|  | n = 33 | n = 34 | p | n = 29 | n = 31 | p | n = 34 | n = 31 | p |
| V<sub>rest</sub> (mV) | -56.7±1.4 | -59.8±1.5 | 0.07 | -50.7±1.5 | -52.1±1.5 | 0.46 | -51.0±1.6 | -52.0±1.6 | 0.62 |
| R<sub>in</sub> (MΩ) | 155±12 | 187±16 | 0.09 | 91±8 | 84±7 | 0.43 | 184±14 | 127±10 | **0.001** |
| Tau (ms) | 38.6±3.7 | 39.6±3.7 | 0.83 | 15.7±1.5 | 15.5±1.5 | 0.96 | 25.0±2.4 | 20.9±2.1 | 0.15 |
| C<sub>m</sub> (pF) | 243±14 | 230±13 | 0.42 | 172±10 | 190±11 | 0.14 | 146±8 | 171±10 | **0.026** |
| AP thresh. (mV) | -28.0±1.1 | 28.4±1.2 | 0.82 | -24.8±1.2 | -27.1±1.2 | 0.13 | -26.9±1.3 | -25.7±1.3 | 0.49 |
| AP amp. (mV) | 76.8±1.4 | 73.2±1.5 | 0.09 | 68.8±1.4 | 71.6±1.4 | 0.15 | 66.8±1.4 | 67.7±1.5 | 0.66 |
| AP h.w. (ms) | 2.18±0.12 | 2.40±0.14 | 0.09 | 0.89±0.05 | 0.91±0.05 | 0.80 | 1.19±0.06 | 0.99±0.05 | **0.001** |
| Depol. rate (mV/ms) | 169±10 | 173±10 | 0.75 | 223±14 | 243±15 | 0.22 | 177±11 | 185±12 | 0.50 |
| Repol. rate (mV/ms) | 32.5±2.0 | 32.6±2.0 | 0.95 | 88.4±5.6 | 91.3±8.7 | 0.64 | 61.9±3.7 | 75.1±4.7 | **0.005** |
| Rheobase (pA) | 151±29 | 142±31 | 0.81 | 364±29 | 464±28 | **0.007** | 189±27 | 350±29 | **0.00014** |
| AHP (mV) | 12.1±0.5 | 12.2±0.7 | 0.92 | 26.0±0.8 | 26.6±0.9 | 0.34 | 21.2±0.5 | 24.7±1.9 | **0.025** |
| H.w. adap. | n.d | n.d |  | 1.11±0.01 | 1.10±0.02 | 0.55 | 1.53±0.06 | 1.47±0.08 | 0.23 |
| AP rate (Hz) | 26.3±3.6 | 23.2±3.1 | 0.21 | 78.9±4.2 | 80.6±8.8 | 0.71 | 36.0±4.7 | 34.4±4.6 | 0.63 |
| Frequency adaptation | 2.17±0.15 | 2.28±0.15 | 0.47 | 1.35±0.09 | 1.34±0.05 | 0.88 | 2.07±0.14 | 1.82±0.13 | 0.09 |

For an explanation of the parameters, see Methods. Thresh. = threshold, amp. = amplitude, h.w. = half-width, depol. = depolarization, repol. = repolarization, AHP = afterhyperpolarization, n.d. = not determined. Values shown are estimated marginal means ± the standard error as determined by implementing a Generalized Linear Mixed Model. P values less than 0.05 are in bold type. The n values are number of neurons obtained from 16 mice (8 WT, 8 *Kcnt1*<sup>m/m</sup>) from 7 litters.

**Table S3.** Electrophysiological parameters of current clamp recordings from acute slices.

|  | Glutamatergic Neurons |  |  | non-Fast Spiking GABA |  |  |
| --- | --- | --- | --- | --- | --- | --- |
|  | WT | Kcnt1<sup>m/m</sup> |  | WT | Kcnt1<sup>m/m</sup> |  |
|  | n = 13 | n = 12 | p | n = 14 | n = 13 | p |
| V<sub>rest</sub> (mV) | -65.5±2.4 | -63.7±2.4 | 0.60 | -60.9±2.2 | -65.5±2.4 | 0.16 |
| R<sub>in</sub> (MΩ) | 124±26 | 106±24 | 0.38 | 299±59 | 161±37 | **0.013** |
| Tau (ms) | 21.3±2.3 | 14.3±2.4 | **0.04** | 18.6±2.3 | 14.5±2.3 | 0.20 |
| C<sub>m</sub> (pF) | 163.9±28 | 133.6±24 | 0.28 | 66.1±11 | 105.8±19 | **0.032** |
| AP thresh. (mV) | -32.6±1.8 | -25±1.9 | **0.006** | -32.1±1.6 | -31.5±1.9 | 0.82 |
| AP amp. (mV) | 77.5±4.1 | 74.2±4.6 | 0.59 | 65.3±4.3 | 75.2±5.1 | 0.08 |
| AP h.w. (ms) | 2.29±0.14 | 2.52±0.16 | 0.30 | 1.26±0.07 | 1.33±0.08 | 0.49 |
| Rheobase (pA) | 158.5±23 | 206.7±24 | 0.16 | 94±22 | 228±23 | **0.003** |
| AHP (mV) | 7.2±1.3 | 7.4±1.2 | 0.91 | 9.8±1.2 | 8.8±1.4 | 0.57 |
| H.w. adaptation | 1.69±0.08 | 1.64±0.08 | 0.64 | 1.33±0.08 | 1.33±0.08 | 0.98 |
| AP rate | 22.9±2.5 | 22.3±2.6 | 0.87 | 37.1±2.5 | 35.2±2.9 | 0.63 |
| Freq. adaptation | 1.71±0.42 | 1.84±0.77 | 0.88 | 3.64±0.56 | 4.41±0.77 | 0.15 |

For an explanation of the parameters, see Methods. Thresh. = threshold, amp. = amplitude, h.w. = half-width. Values shown are estimated marginal means ± the standard error as determined by implementing a Generalized Linear Mixed Model. P values less than 0.05 are in bold type. The n values are number of neurons obtained from 16 mice (9 WT, 7 Kcnt1<sup>m/m</sup>) from 8 litters.

**Table S4.** Synaptic current data from neuronal cultures.

|  | Excitatory neurons |  |  | Inhibitory neurons |  |  |
| --- | --- | --- | --- | --- | --- | --- |
|  | WT | Kcnt1<sup>m/m</sup> |  | WT | Kcnt1<sup>m/m</sup> |  |
|  | n = 56 | n = 57 | p | n = 37 | n = 45 | p |
| sEPSC freq. (Hz) | 3.0±0.86 | 5.6±0.76 | **0.001** | 10.4±2.0 | 11.4±2.0 | 0.93 |
| sEPSC amp. (pA) | 17.6±1.2 | 20.1±1.4 | 0.16 | 30.8±2.2 | 32.2±2.0 | 0.60 |
| sEPSC decay (ms) | 3.5±0.07 | 3.5±0.07 | 0.83 | 2.7±0.08 | 2.9±0.09 | 0.19 |
|  | n = 28 | n = 25 |  | n = 21 | n = 26 |  |
| sIPSC freq. (Hz) | 2.1±0.3 | 2.7±0.4 | 0.09 | 2.6±0.4 | 2.9±0.4 | 0.20 |
| sIPSC amp. (pA) | 27.4±2.3 | 32.5±2.5 | 0.17 | 25.8±2.5 | 27.6±2.4 | 0.63 |
| sIPSC decay (ms) | 18.4±1.3 | 19.5±1.3 | 0.17 | 16.5±1.4 | 17.0±1.3 | 0.57 |
| E/I ratio | 0.34±0.05 | 0.45±0.06 | **0.037** | 0.74±0.09 | 0.66±0.08 | 0.36 |

For an explanation of the parameters, see Methods. amp. = amplitude, freq. = frequency, E/I = excitation inhibition. Values shown are estimated marginal means ± the standard error as determined by implementing a Generalized Linear Mixed Model. P values less than 0.05 are in bold type. The n values are number of neurons obtained from 4 litters and 8 mice (4 WT, 4 *Kcnt1<sup>m/m</sup>*). \*GABAergic neurons were not subclassified for sIPSC measurements because the internal solution to measure sIPSCs alters many AP parameters.

**Figure S1.**
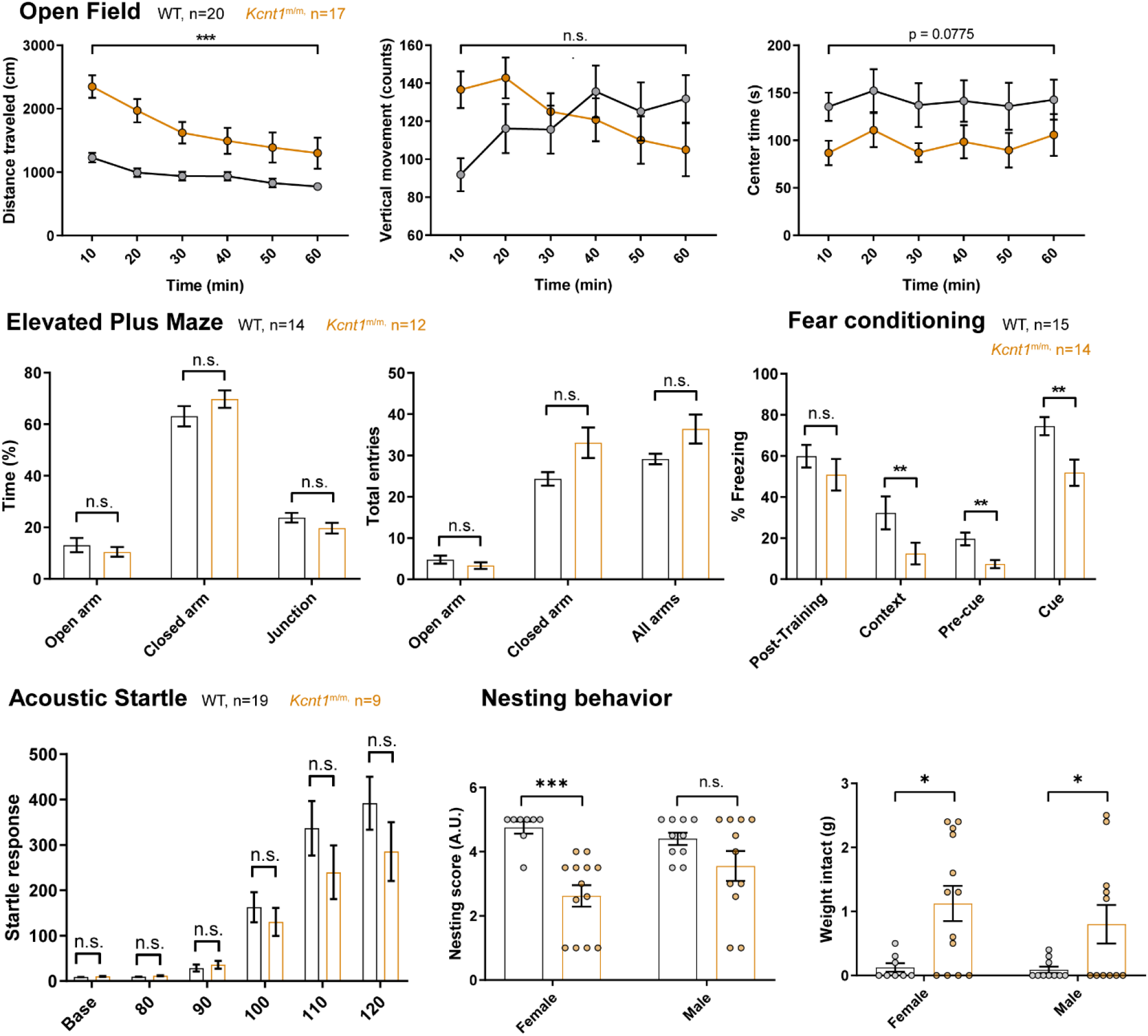
Behavioral phenotypes in *Kcnt1*^m/m^ mice. **(A)** Open field test: distance traveled, vertical exploration and center time. ***p<0.001; Two-way repeated measures ANOVA. **(B)** Elevated Plus Maze test assessing % of time spent in open arms, closed arms or at the junction (left) and total number of entries in the arms (right). Unpaired two-tailed t-test with Holm-Sidak correction. **(C)** Fear conditioning test for cued and contextual memory. **p<0.01; Mann-Whitney U test. **(D)** Acoustic Startle Response. Two-way repeated measures ANOVA. **(E)** Nesting behavior test assessing nest quality (left; nesting score) and weight of remaining intact nesting material (right). n = 8 WT female, 10 WT males, 13 *Kcnt1*^m/m^ females, and 11 *Kcnt1*^m/m^ males. *p<0.05; ***p<0.001; Unpaired two-tailed t-test with Holm-Sidak correction. n.s., non significant.

**Supp Fig 2.**
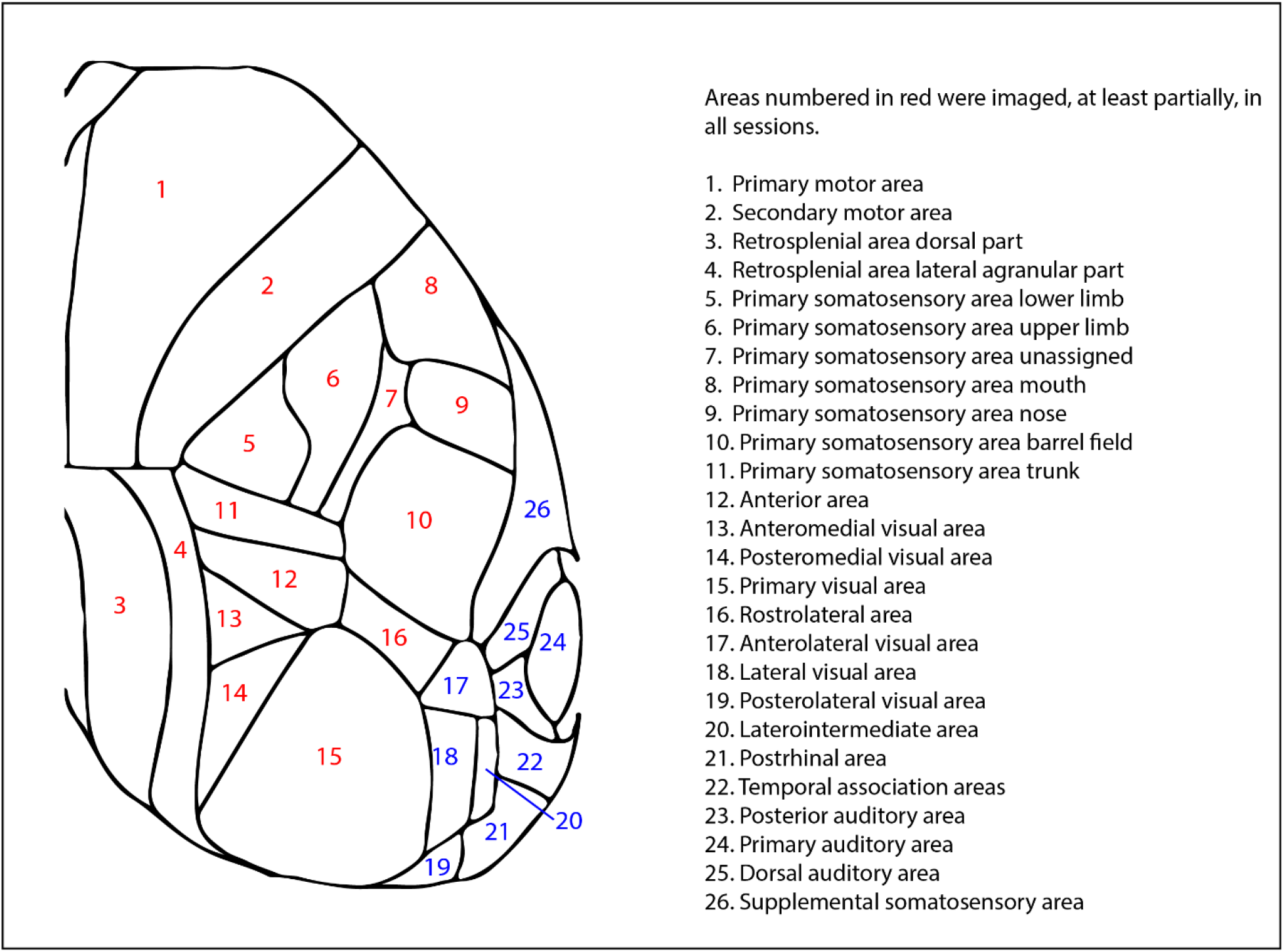
Allen CCF Cortical Area Names

**Figure S3.**
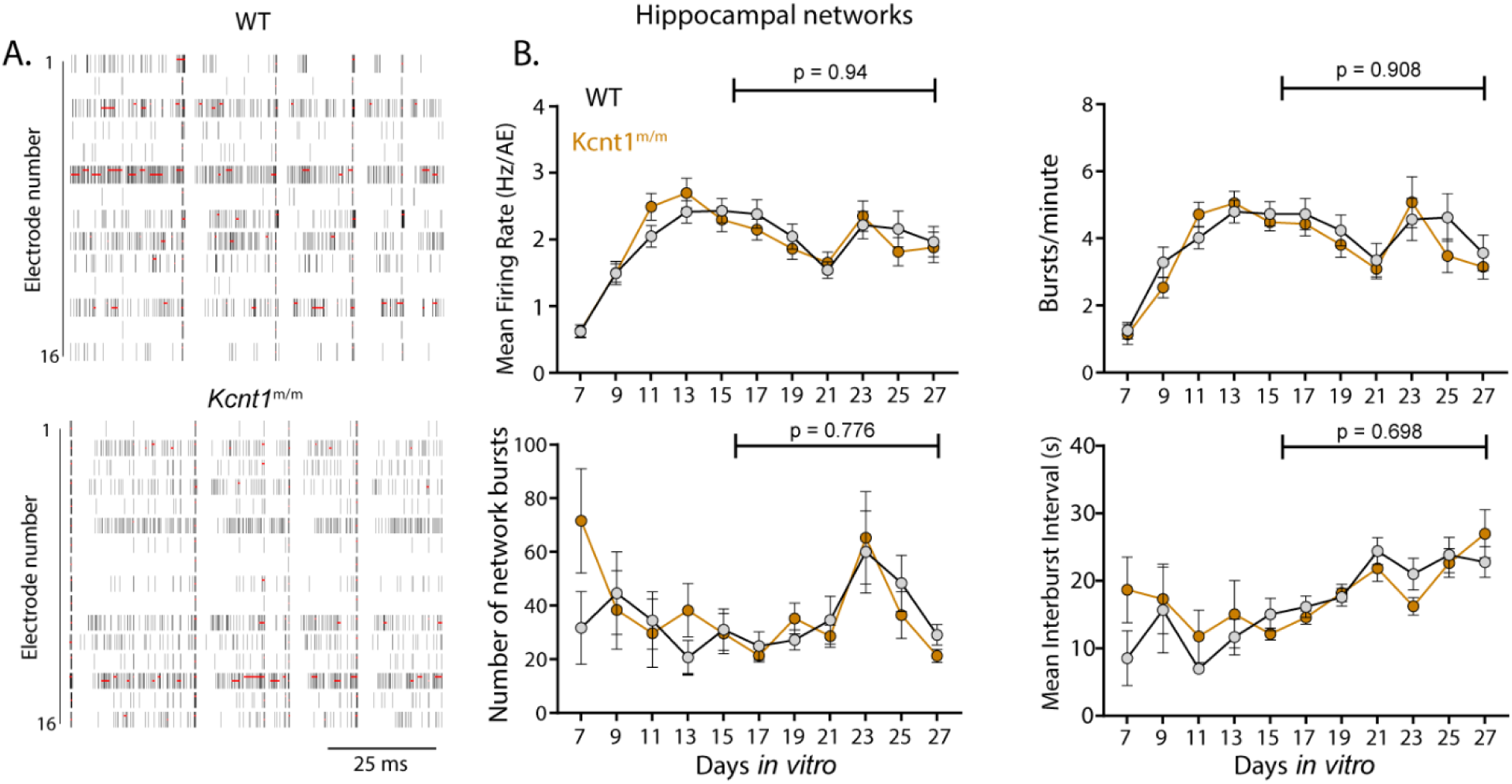
*In vitro* hippocampal networks from *Kcnt1*^m/m^ mice show normal population activity. **(C)** Raster plots showing WT and *Kcnt1*^m/m^ hippocampal network firing across the 16 electrodes of a well of a representative plate at DIV 21. The black bars indicate spikes, and the red bars indicate bursts. **(D)** Graphs plotting measures of spontaneous activity as a function of days *in vitro* on MEAs of WT (gray, n=53 wells, 5 mice) and *Kcnt1*^m/m^ (orange, n=53 wells, 5 mice) hippocampal neurons. MFR per active electrodes, number of bursts per minute, number of network bursts, and interburst interval are shown. Permutated p-values for mature DIV 17 to 27 calculated with a Mann-Whitney U test followed by 1,000 permutations are indicated on the graphs.

